# Spatial immune ecosystems govern therapeutic response in HER2-low breast cancer

**DOI:** 10.64898/2026.08.19.745800

**Authors:** Olajumoke Ogunlusi, Soupayan Banerjee, Akshara R. Singareeka, Stephen Akanbi, Mrinmoy Sarkar, Pritam Dey, Bo Lin, Yingxiang Xu, Tommy Tran, Danielle Fails, Bani Mallick, Gabriela Raso, Debasish Tripathy, Tapasree Roy Sarkar

## Abstract

HER2-low breast cancer represents a clinically important but biologically heterogeneous disease state, and the spatial immune programs underlying therapeutic response remain poorly understood. Here, we used single-cell spatial transcriptomics to characterize HER2-low and HER2-high breast tumors and define microenvironmental features associated with treatment sensitivity and resistance. We identified diverse malignant, stromal, and immune compartments, with dendritic cells emerging as a highly remodeled population in HER2-low tumors. Focused analysis resolved distinct dendritic-cell states, including homeostatic cDC2, IFN-activated mature cDC, classical functional cDC2, plasmacytoid DC, and ITGAX-positive monocyte-derived DC populations. Spatial proximity analysis further revealed that resistant HER2-low tumors exhibited increased segregation of tumor epithelial cells from effector immune populations and enrichment of myeloid-rich immune niches, consistent with an immune-restricted spatial architecture. Independent TCGA-BRCA validation confirmed the clinical relevance of these dendritic-cell states, with elevated homeostatic cDC2 signatures predicting poor survival, whereas inflammatory dendritic-cell signatures were associated with favorable outcomes. Resistant HER2-low tumors were characterized by enrichment of homeostatic and classical cDCs, depletion of IFN-activated cDCs and pDCs, altered tumor–myeloid–T-cell communication, and expansion of spatially organized resistant niches, whereas sensitive tumors retained immune-intermixed niches enriched for antigen presentation and effector immune interactions. Together, these findings demonstrate that therapeutic resistance in HER2-low breast cancer is driven by coordinated spatial remodeling of dendritic-cell states and immune architecture, identifying dendritic-cell–myeloid niche organization as a potential biomarker and therapeutic vulnerability.

## Introduction

Breast cancer is a biologically heterogeneous disease whose clinical behavior is governed not only by tumor-intrinsic genetic alterations but also by spatially organized interactions with the surrounding tumor microenvironment (TME)^1^. Bulk transcriptomic analyses obscure this heterogeneity by averaging signals across malignant, stromal, and immune compartments, while single-cell RNA sequencing (scRNA-seq), although transformative, disrupts tissue architecture and eliminates information about cellular localization and neighborhood relationships^2^. Consequently, how transcriptional programs are spatially deployed within intact breast tumors—particularly across disease stages and molecular subtypes—remains poorly understood. Spatial transcriptomics has emerged as a powerful approach to overcome these limitations by enabling transcriptome-wide gene expression profiling while preserving histological context^3^. By retaining spatial information, these technologies have revealed functionally distinct cellular niches, spatially restricted immune states, and microenvironmental signaling gradients in multiple solid tumors^4^. Recent advances have further increased resolution and compatibility with formalin-fixed paraffin-embedded (FFPE) tissues, allowing the analysis of clinically annotated human tumor specimens at near–single-cell scale^5^. Despite these advances, the application of spatial transcriptomics to breast cancer remains relatively limited compared with other tumor types. Existing studies have largely focused on specific contexts, such as immune infiltration patterns or stromal organization in small cohorts, often without stratification by disease stage or molecular subtype^6^. As a result, the spatial principles governing breast tumor evolution—from early lesions to advanced disease—remain incompletely defined.

This gap is particularly evident for HER2-low breast cancer, a recently recognized clinical entity characterized by low but detectable HER2 expression that does not meet criteria for HER2 amplification^7,8^ HER2- low tumors constitute a substantial fraction of both hormone receptor–positive and triple-negative breast cancers and exhibit distinct biological behavior and therapeutic vulnerabilities, including responsiveness to next-generation HER2-targeted antibody–drug conjugates^9^. This heterogeneity suggests that treatment response may not be determined solely by HER2 expression level or tumor-cell-intrinsic programs, but also by the broader tumor microenvironment. The tumor microenvironment is a complex multicellular ecosystem composed of cancer cells, stromal cells, immune cells, endothelial cells, extracellular matrix, and soluble signaling factors^10^. These non-malignant components actively regulate tumor progression, immune surveillance, metastatic potential, and therapeutic response^10^. In breast cancer, the spatial organization of tumor cells with stromal and immune populations can strongly influence whether a tumor remains immune-accessible or becomes immune-excluded and therapy-resistant^11^. Therefore, defining how specific microenvironmental cell types are organized within sensitive versus resistant HER2-low tumors may reveal mechanisms of treatment response and identify new biomarkers or therapeutic vulnerabilities.

Dendritic cells (DCs) are central regulators of anti-tumor immunity because they capture tumor antigens, process and present antigens to T cells, and coordinate adaptive immune responses^12,13^. Functionally active DCs can promote cytotoxic T-cell priming and support immune-mediated tumor control, particularly after chemotherapy-induced tumor-cell death^12^. However, DC populations within tumors are heterogeneous and can exist in homeostatic, inflammatory, interferon-activated, tolerogenic, or myeloid-derived states^13^. In a suppressive tumor microenvironment, DCs may become dysfunctional, spatially excluded from tumor regions, or unable to effectively activate T cells^14^. Therefore, the abundance, functional state, and spatial localization of DC populations may critically influence whether chemotherapy promotes productive anti-tumor immunity or fails to overcome immune suppression.

Here, we analyzed a retrospective cohort of 21 HER2-expressing breast tumor (HER2-high and HER2- low) specimens collected over the last 10 years, including tumors from patients treated with neoadjuvant therapy and annotated for pathologic response to treatment. We focused on chemotherapy-sensitive (achieving pathologic complete response or pCR to pre-operative chemotherapy) versus chemotherapy-resistant (non-pCR) (**Supplementary Table 1)** HER2-low tumors to determine how treatment response is associated with tumor microenvironmental organization. To place these findings in a broader HER2-context, we also compared the spatial architecture of HER2-low tumors with HER2-high sensitive and resistant tumors. This comparative framework allowed us to distinguish HER2-low-specific resistance features from more general mechanisms of therapy resistance across HER2-defined breast cancer states. We focused on the spatial relationships among dendritic-cell populations along with other immune cells, and tumor-cell states to determine whether resistant tumors exhibit distinct microenvironmental organization. Our analysis identifies response-associated spatial niches and reveals that chemotherapy-resistant HER2-low tumors are characterized by a distinct remodeling of dendritic-cell states, including increased abundance of homeostatic cDC2 and classical functional cDC2 populations, together with reduced IFN-activated mature cDC and plasmacytoid DC (pDC) populations. This altered DC landscape was accompanied by CAF-rich stromal expansion, reduced cytotoxic immune infiltration, immune-suppressed signaling, and spatial compartmentalization of the tumor microenvironment. Together, these findings support a model in which impaired DC activation, CAF-driven stromal remodeling, and immune exclusion cooperate to weaken antitumor immunity and promote therapeutic resistance in HER2-low breast cancer.

## Results

### Study design and single-cell-resolution spatial atlas

To investigate the molecular determinants of therapy response and the cellular architecture of the tumor ecosystem in HER2-low breast cancer, we applied spatially resolved molecular profiling to interrogate gene expression programs, cellular states, and multicellular organization within their native tissue context. Spatial omics analyses were performed on a retrospective cohort of 21 human breast tumor specimens, including HER2-low (n=16) and HER2-high tumors (n=5) who underwent surgical resection with (n=20) or without (n=1) neoadjuvant treatment **(Figure 1A, supplementary table 1**). The cohort included both therapy-sensitive and therapy-resistant tumors, allowing direct comparison of spatial features associated with effective treatment response versus residual resistant disease.

**Figure 1.**
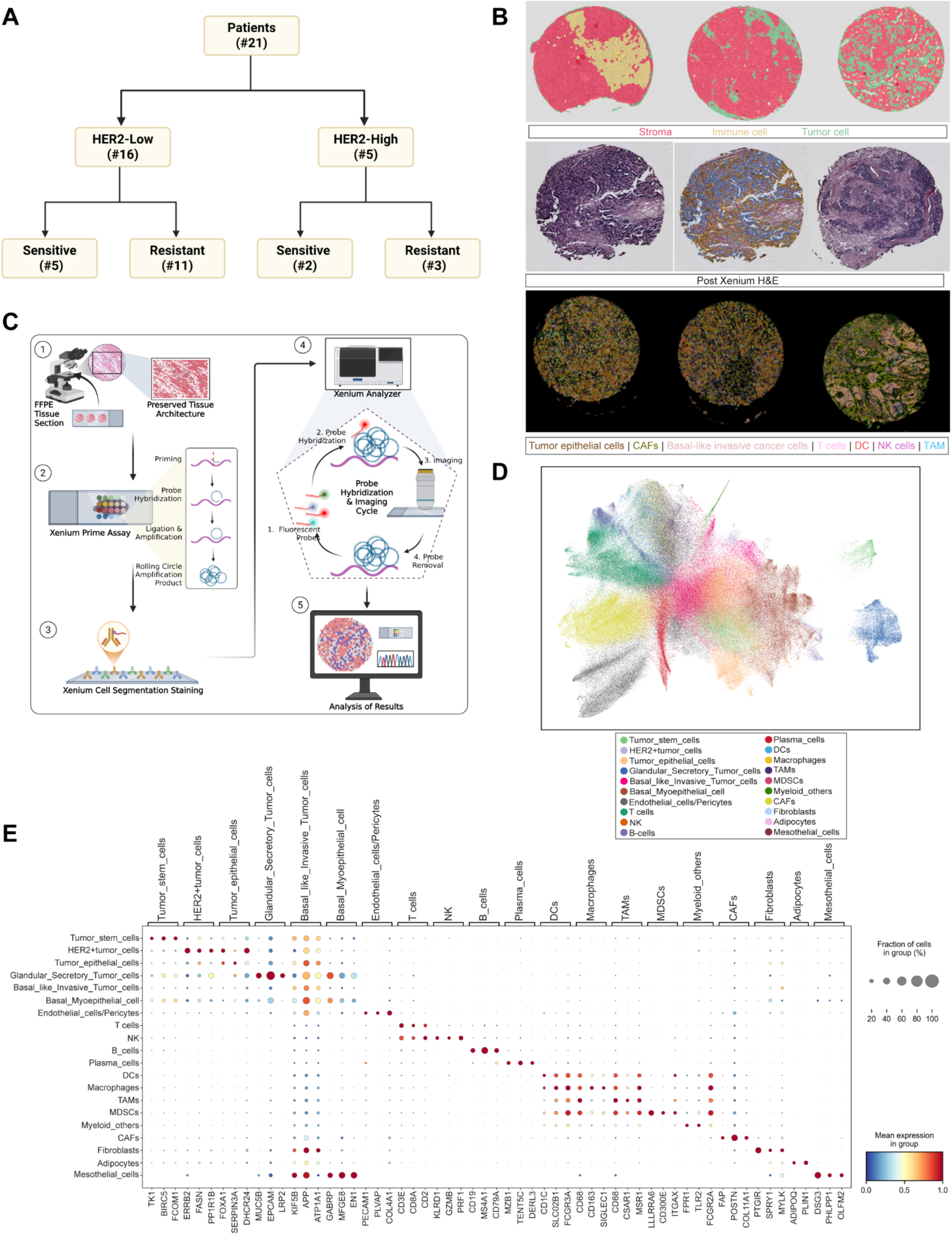
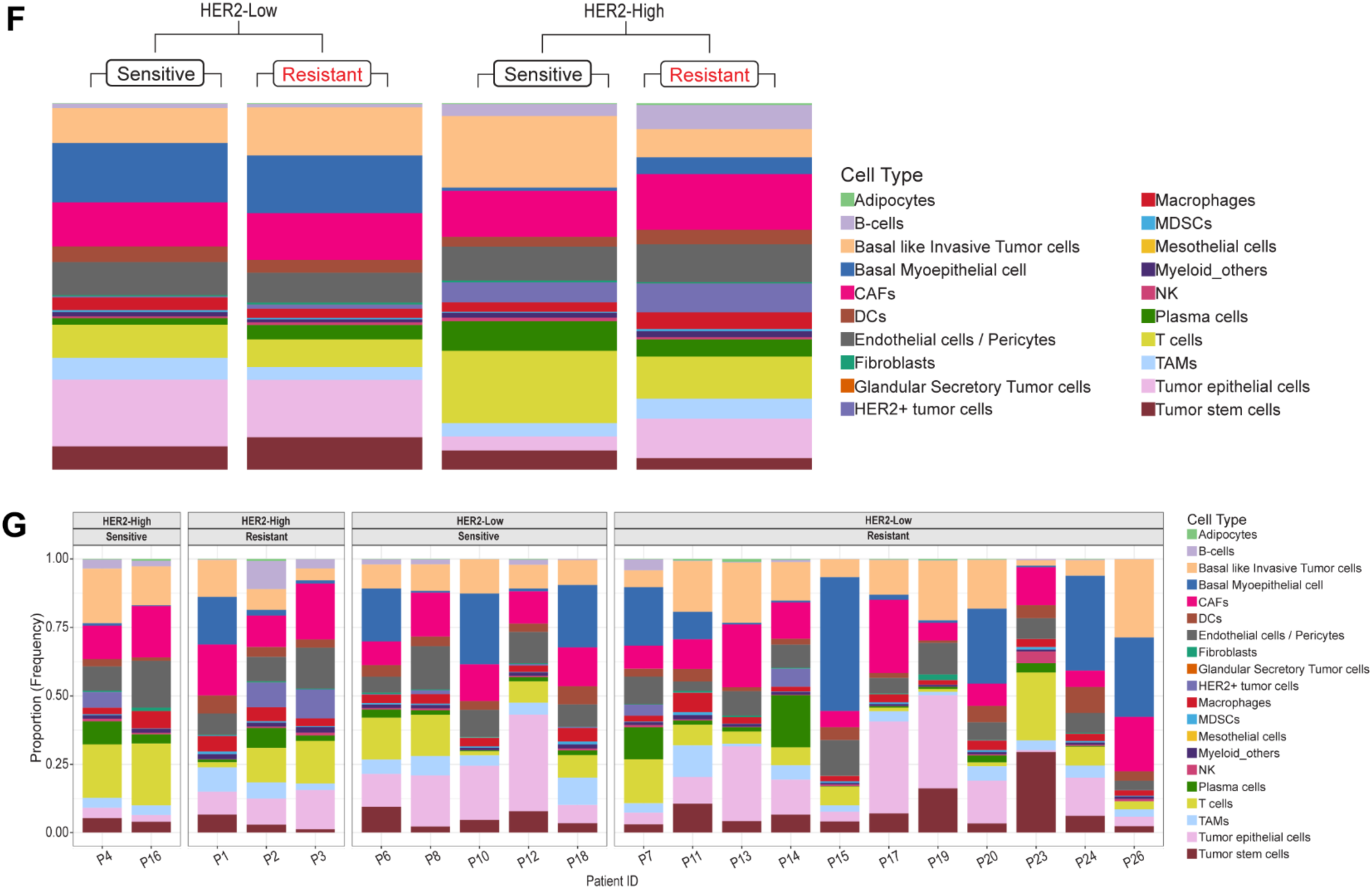
Single-cell spatial transcriptomic profiling reveals the cellular landscape of HER2-low and HER2-high breast tumors and subtype-specific cellular composition associated with therapeutic response. (A) Study design and patient cohort. Twenty-one breast tumor specimens were analyzed by single- cell spatial transcriptomics and stratified according to HER2 status and therapeutic response. The cohort comprised 16 HER2-low tumors (5 sensitive [achieving a pathological complete response, pCR] and 11 resistant, (not achieving pCR)] and 5 HER2-high tumors (2 sensitive and 3 resistant). (B) Representative images of breast tumor sections showing digital pathology (upper panel), post xenium H&E staining (middle panel), spatial transcriptomic segmentation, and reconstructed cellular architecture following Xenium profiling (lower panel). (C) Experimental workflow for Xenium-based spatial transcriptomic analysis. Formalin-fixed paraffin-embedded (FFPE) tumor sections were processed using the Xenium *in Situ* platform, including tissue preparation, probe hybridization, rolling-circle amplification, fluorescence imaging, cell segmentation, transcript decoding, and downstream single-cell spatial analysis. (D) UMAP visualization of integrated single-cell spatial transcriptomic data identifying major cellular populations across all tumors. Unsupervised clustering resolved diverse tumor, stromal, vascular, and immune cell populations, including tumor epithelial cells, fibroblasts, endothelial cells, myeloid cells, lymphocytes, plasma cells, adipocytes, and other stromal components. (E) Dot plot showing canonical marker gene expression across all annotated cell populations. Dot size represents the proportion of cells expressing each marker, whereas color intensity indicates the average normalized expression level, confirming robust annotation of individual cell types. (F) Comparison of overall cellular composition between sensitive and resistant tumors within HER2-low and HER2-high breast cancer. Stacked bar plots summarize the relative abundance of major cell populations, demonstrating subtype-specific remodeling of the tumor microenvironment associated with therapeutic response. (G) Patient-level cellular composition across individual HER2-low and HER2-high tumors. Each stacked bar represents one patient, illustrating inter-patient heterogeneity while highlighting consistent differences in cellular composition between sensitive and resistant tumors.

Archived formalin-fixed, paraffin-embedded (FFPE) tumor specimens were used, consistent with current clinical pathology standards for tissue processing and long-term preservation^15^. We selected FFPE sections that contained histologically defined tumor regions, stromal compartments, and residual tumor areas. These regions included areas with transcriptomic and histologic features of malignant tumor cells, surrounding stroma, immune-enriched regions, and invasive tumor boundaries, which are critical for understanding tumor-cell and microenvironmental characteristics in the tumor ecosystem. The images from digital pathology, post Xenium H&E and spatial transcriptomics are shown in **Figure 1B**. These annotations were integrated with spatial transcriptomic profiles to determine how malignant epithelial cells, CAFs, immune cells, endothelial cells, and extracellular matrix-associated programs were spatially organized across therapy-sensitive and therapy-resistant tumors. This study design enabled us to compare the spatial architecture of HER2-low sensitive versus HER2-low resistant tumors and to determine whether treatment resistance is associated with distinct tumor microenvironmental states, including immune exclusion, CAF enrichment, stromal remodeling, altered tumor– immune proximity, and residual tumor-cell survival niches. In parallel, comparison with HER2-high sensitive and resistant tumors allowed us to determine whether spatial resistance programs are shared across HER2-defined breast cancer states or are uniquely enriched in HER2-low disease.

Spatial transcriptomic profiling was performed using the 10x Genomics Xenium in Situ platform, which enables highly multiplexed, single-molecule RNA detection within intact tissue sections while preserving spatial context. The overall workflow for this study is depicted in **Figure 1C**. The xenium spatial transcriptome sequencing^3^ performed with FFPE tissue sections (5–10 μm) was mounted onto Xenium-compatible slides, followed by fixation, permeabilization, and probe hybridization according to the manufacturer’s protocol (method). Using this approach, we sought to identify generalized factors that govern cancer cell fate. To validate the 5,000-plex Xenium human breast panel, we evaluated gene expression across expected cell types and transferred scFFPE-seq annotations using supervised labeling. This approach enabled robust identification of diverse cellular populations, confirming the capture of tumor heterogeneity.

### HER2-low sensitive and resistant tumors display distinct TME composition

Xenium successfully identified basal myoepithelial cells, CTA^+^ tumor stem cells^16^, invasive tumor cells, mast cells, lymphocytes and myeloid cells, fibroblasts, and CAFs that comprise the TME of HER2-low and HER2-high breast tumors. The dimensionality reduction separated epithelial, immune, and stromal populations into distinct transcriptional compartments by UMAP (**Figure 1D)**, revealing substantial cellular heterogeneity across the cohort. Major annotated populations included tumor stem-like cells, HER2-positive tumor cells, general tumor epithelial cells, glandular secretory tumor cells, basal-like invasive tumor cells, basal myoepithelial-like cells, endothelial cells/pericyte, T cells, NK cells, B cells, plasma cells, dendritic cells, macrophages, tumor-associated macrophages, myeloid-derived suppressor cells, additional myeloid populations, CAFs, fibroblasts, adipocytes, and mesothelial cells. Distinct tumor-cell states included proliferative/stem-like tumor cells, HER2-enriched tumor cells, glandular secretory tumor cells, basal-like invasive tumor cells, and basal myoepithelial-like cells. This organization suggests that treated breast tumors contain multiple malignant programs rather than a single uniform tumor-cell state. T cells, NK cells, B cells, and plasma cells occupied distinct regions, supporting the ability to resolve adaptive and innate immune populations. Whereas myeloid cells such as dendritic cells, macrophages, TAMs, MDSCs, and other myeloid populations, formed a separate transcriptional compartment, consistent with diverse inflammatory and immunosuppressive myeloid states within the tumor microenvironment. Stromal populations, including CAFs, fibroblasts, endothelial cells/pericytes, and adipocytes, were also clearly distinguished, supporting robust identification of nonmalignant structural and vascular components of the tumor ecosystem. Marker-dot plot analysis (**Figure 1E)** validated the cell-type annotations^17–36^. The stacked bar plots show that cellular composition varies substantially between tumors stratified by treatment response (sensitive vs resistant) (**Figure 1F, supplementary Figure 1)**, as well as across all the HER2-low and HER2-high patients **(Figure 1G)**, indicating marked inter-patient heterogeneity in both the tumor-cell compartment and the tumor microenvironment. Across the cohort (**Figure 1G**), each tumor contains a unique mixture of tumor epithelial populations, immune cells, and stromal cells, suggesting that response status is associated not with a single cell type alone, but with the broader balance of these compartments. Together, these data establish a high-resolution cellular atlas of HER2-low and HER2-high breast tumors following neoadjuvant therapy. This cellular framework enables investigation of whether resistant HER2-low tumors are associated with specific tumor-cell states, immune-suppressive myeloid populations, CAF-enriched niches, or altered tumor–stromal–immune organization within the tumor microenvironment.

### DC-state remodeling distinguishes treatment response

Because dendritic cells are central regulators of antigen presentation, T-cell priming, and antitumor immunity, we next re-clustered the dendritic-cell compartment to determine whether specific DC states were associated with treatment sensitivity or resistance. The relevance of DCs to neoadjuvant response is further supported by emerging clinical studies using DC-based strategies in breast cancer^37^. Intratumoral cDC1 therapy and HER2-directed DC vaccination have shown feasibility, immune activation, and preliminary evidence of enhanced anti-tumor responses when combined with HER2-targeted or neoadjuvant therapy^38^. The UMAP projection demonstrated clear separation of five different DC states (**Figure 2A).** Unsupervised re-clustering resolved five transcriptionally distinct DC populations in our studied tumors: classical functional cDC2^39^, homeostatic cDC2^40^, IFN-activated mature cDC^39^, ITGAX^+^ monocyte-derived DC^39^, and plasmacytoid DCs (pDCs)^39^.

**Figure 2.**
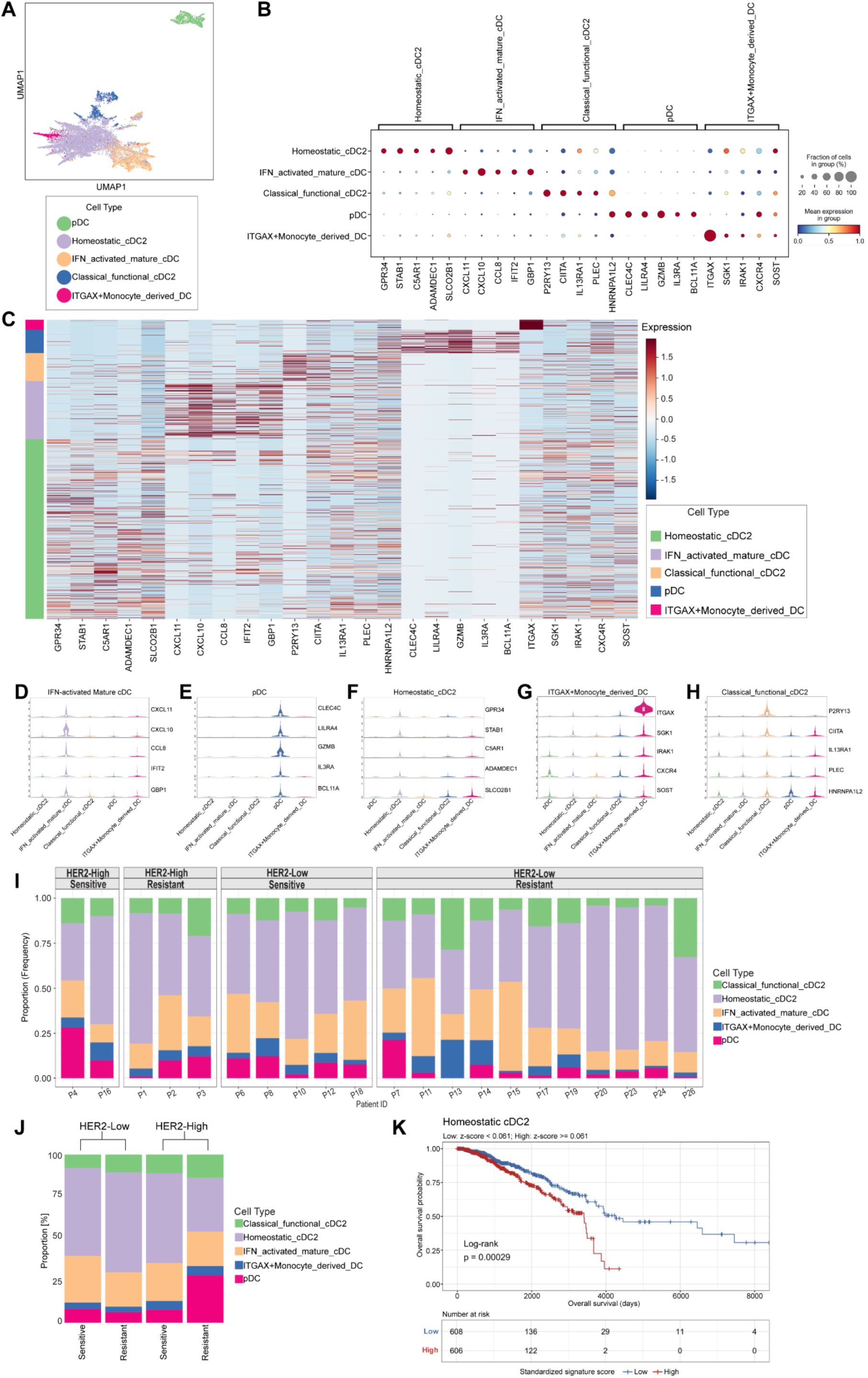
Single-cell characterization of dendritic-cell heterogeneity reveals subtype-specific remodeling associated with therapeutic response. **(A)** UMAP visualization of dendritic cells identified from HER2 breast tumors, revealing five transcriptionally distinct populations: plasmacytoid dendritic cells (pDCs), homeostatic cDC2, IFN-activated mature cDCs, classical functional cDC2, and ITGAX^+^ monocyte-derived dendritic cells (ITGAX⁺ MoDCs). **(B)** Dot plot showing the expression of canonical marker genes used to define each dendritic- cell population. Dot size represents the percentage of cells expressing each gene, and color intensity indicates the average normalized expression level. **(C)** Heatmap of differentially expressed genes across the five dendritic-cell subsets, highlighting the distinct transcriptional programs that define each population. Columns represent genes and rows represent individual cells grouped by dendritic-cell subtype. Violin plots illustrating the expression distribution of representative marker genes defining each dendritic-cell subset **(D)** IFN-activated mature cDCs, **(E)** pDCs, **(F)** homeostatic cDC2, **(G)** ITGAX^+^ monocyte-derived DCs, and **(H)** classical functional cDC2. These subtype-specific marker profiles further validate the transcriptional identity of each dendritic-cell population. **(I)** Relative abundance of dendritic-cell populations in individual HER2-high and HER2-low tumors stratified by therapeutic response (sensitive versus resistant). Each bar represents one patient, illustrating inter- patient heterogeneity in dendritic-cell composition. **(J)** Summary of dendritic-cell composition across HER2-high and HER2-low tumors. **(K)** Kaplan–Meier survival analysis of the TCGA-BRCA cohort (n = 1,214) stratified by dendritic-cell (DC) signature scores. Patients were dichotomized into high- and low-expression groups based on the median signature score derived from standardized (z-score) expression of marker genes for homeostatic cDC2 (STAB1, IL13RA1, SLC02B1).

Homeostatic cDC2 represented the major transcriptional backbone of the DC compartment, whereas IFN-activated mature cDCs and classical functional cDC2 occupied more discrete regions, suggesting functionally specialized antigen-presenting states. pDCs formed a distinct cluster, consistent with their unique transcriptional identity. In contrast, ITGAX^+^ monocyte-derived DCs localized to a separate branch-like region, suggesting an inflammatory or monocyte-derived differentiation state within the tumor microenvironment. Marker analysis (**Figure 2B**) confirmed the identity of these DC subsets, pDCs were enriched for CLEC4C, LILRA4, and IL3RA, supporting their plasmacytoid identity^41^. Homeostatic cDC2 showed enrichment of STAB1, consistent with a steady-state or tissue-resident DC-like program^42^ along with SLCO2B1^43^. IFN-activated mature cDCs expressed interferon- and maturation-associated genes, including CXCL10, CXCL11, and IFIT2, indicating an activated antigen-presenting state^44^. Classical functional cDC2 showed expression of P2RY13^45^ and CIITA^46^, consistent with antigen-processing and MHC-II-associated function. ITGAX-positive monocyte-derived DCs were marked by strong ITGAX expression^47^, supporting their assignment as monocyte-derived inflammatory DC-like cells. Marker analysis (**Figures 1E and 2B, C**) confirmed that, along with established DC markers, the identified DC subsets exhibited minimal expression of macrophage-associated markers, including SIGLEC1^48^, CD163 and MSR1^49^ further supporting their classification as bona fide dendritic-cell populations rather than macrophages.

The accompanying heatmap (**Figure 2C)** further confirmed lineage-specific transcriptional programs across the different types of DC cells. The violin plots for each DC population show the expression distribution of specific marker genes across the different DC subsets **(Figure 2D-H)**. We next examined the distribution of DC states across all the HER2-low and HER2-high patients (**Figure 2I**) as well as tumors stratified by treatment response (**Figure 2I-J**). The proportional bar plot showed that DC composition varied substantially across clinical groups. In HER2-low tumors, treatment sensitivity and resistance appear to be associated with distinct dendritic-cell states rather than a simple increase or loss of total DCs. Resistant tumors show higher homeostatic cDC2 and classical functional cDC2, but reduced IFN-activated DCs and reduced pDCs. This suggests that resistant tumors may retain antigen-presenting DC populations, but these DCs may not be fully activated into an interferon-driven inflammatory state. In other words, resistant tumors may contain DCs that are present and potentially antigen-processing competent, but less immunostimulatory. The reduction of IFN-activated DCs suggests that resistant HER2-low tumors may have impaired immune activation, even if DCs are still present. This could contribute to poor CD8 T-cell priming, weaker NK/T-cell recruitment, and ineffective antitumor immunity after therapy^50^. By contrast, sensitive HER2-low tumors show higher pDCs and higher ITGAX^+^ monocyte-derived DCs. This suggests that sensitive tumors may have a more inflammatory or therapy-responsive myeloid/DC environment. pDCs can produce type I interferon under certain conditions^51^, and ITGAX^+^ monocyte-derived DCs may reflect therapy-induced inflammatory recruitment or antigen-processing activity^47,52^. In sensitive tumors, these populations may contribute to immune activation, tumor antigen sensing, and recruitment of effector immune cells.

In contrast, HER2-high resistant tumors displayed marked remodeling of the DC compartment (**Figure 2I-J)**, characterized by a pronounced expansion of pDCs and a substantial reduction in Homeostatic cDC2 relative to HER2-high sensitive tumors. Classical functional cDC2 and ITGAX⁺ monocyte-derived DCs also showed modest increases in HER2-high resistant tumors. These results indicate that resistance is associated with distinct DC states depending on HER2 expression, with a Homeostatic cDC2-enriched phenotype in HER2- low tumors and a pDC-enriched phenotype in HER2-high tumors. Analysis of the TCGA-BRCA cohort independently validated our spatial observations, demonstrating that a high homeostatic cDC2 signature was associated with poor overall survival (log-rank p = 0.00029; FDR-adjusted p ≈ 0.0014) (**Figure 2K),** whereas elevated ITGAX⁺ monocyte-derived DC and IFN-activated mature cDC signatures predicted favorable survival **(Supplementary Figure 2)**. No significant prognostic associations were observed for pDC or classical functional cDC2 signatures **(Supplementary Figure 2)**. Together, this study identifies previously unappreciated heterogeneity within the dendritic-cell compartment of treated HER2-low and HER2-high breast tumors.

Gene ontology analysis revealed that each DC state (**Supplementary figure 3)** was associated with a distinct biological program, supporting functional heterogeneity within the dendritic-cell compartment of treated HER2-low and HER2-high breast tumors^52–54^. The KEGG pathway enrichment analysis across distinct DC-related clusters (**Supplementary figure 4)** revealed that dendritic-cell subsets in HER2-low tumors are not functionally uniform, but instead occupy distinct homeostatic, antigen-presenting, inflammatory, interferon-associated, and myeloid-derived states^55–59^. Together, GO enrichment and KEGG analyses suggest that therapeutic resistance is accompanied by coordinated functional and communication remodeling of the tumor microenvironment. Resistant tumors showed enrichment of immune-regulatory, inflammatory, myeloid and stromal-associated programs, together with expanded tumor–immune–myeloid communication networks.

## HER2-low resistant tumors exhibit altered tumor–immune communication networks

CellChat analysis revealed subtype-specific remodeling of tumor–immune communication in resistant tumors (**Figure 3A**). In HER2-low tumors, resistant samples exhibited a marked increase in inferred cell–cell interactions compared with sensitive tumors (3459 vs. 2524), whereas overall interaction strength remained similar (0.052 vs. 0.051) (**Figure 3B; Supplementary Figure 5A**). The resistant HER2-low network (**Figure 3C, D**) showed increased interactions among tumor cells, macrophages, NK cells, CD8 T cells, and multiple dendritic-cell subsets, particularly homeostatic cDC2 and classical functional cDC2 populations. In HER2-high tumors, resistant samples also exhibited increased tumor–immune–myeloid connectivity compared with sensitive tumors (**Figure 3E**), accompanied by higher interaction number and interaction strength (**Figure 3F; Supplementary Figure 5B**). Network visualization demonstrated broader communication among tumor, immune, and myeloid populations in resistant HER2-high tumors (**Figure 3G, H).**

**Figure 3.**
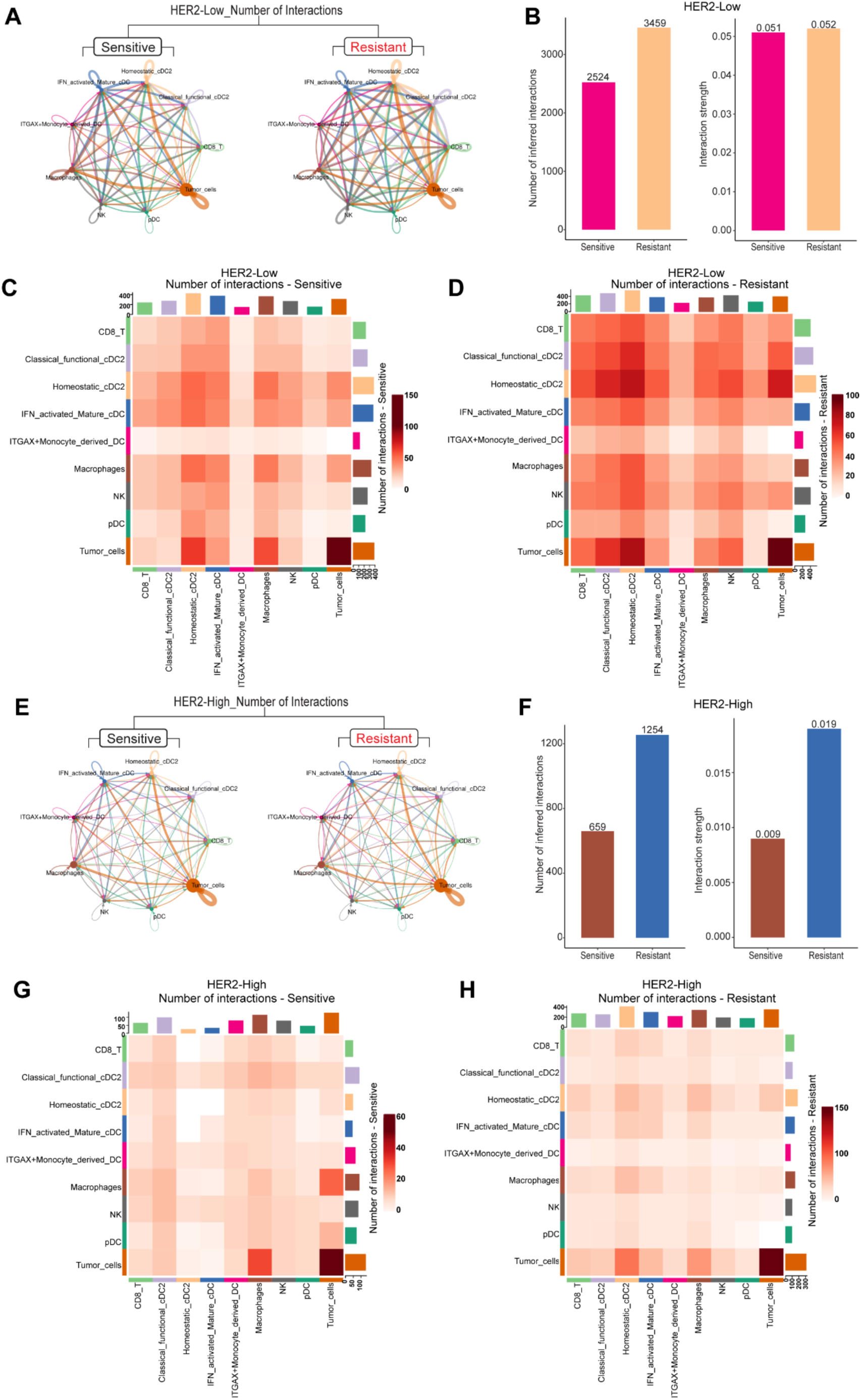
Cell–cell communication networks are extensively remodeled during therapeutic resistance in HER2-low and HER2-high breast tumors. **(A)** Cell–cell interaction networks inferred from spatial transcriptomic data in HER2-low sensitive and HER2-low resistant tumors. Nodes represent major cellular populations, and edge thickness is proportional to the number of predicted ligand–receptor interactions between cell types. **(B)** Quantification of the total number of inferred cell–cell interactions and the overall interaction strength in HER2- low tumors. Resistant tumors exhibit increased numbers of predicted interactions and greater overall interaction strength compared with sensitive tumors, indicating enhanced intercellular communication during therapeutic resistance. **(C, D)** Heatmaps showing the frequency of pairwise cell–cell interactions among major immune and tumor cell populations in HER2-low sensitive **(C)** and HER2-low resistant **(D)** tumors. Resistant tumors display increased interaction frequencies involving tumor cells, dendritic-cell subsets, macrophages, CD8⁺ T cells, and NK cells, consistent with extensive remodeling of the communication network. **(E)** Cell–cell interaction networks in HER2-high sensitive and HER2-high resistant tumors. Compared with sensitive tumors, resistant HER2-high tumors demonstrate a denser interaction network with increased connectivity among immune and tumor cell populations. **(F)** Quantitative comparison of the total number of inferred interactions and overall interaction strength in HER2-high tumors. Resistant tumors exhibit a marked increase in both interaction number and communication strength relative to sensitive tumors, indicating enhanced cellular cross-talk during disease progression. **(G, H)** Heatmaps illustrating pairwise interaction frequencies among major cell populations in HER2-high sensitive **(G)** and HER2-high resistant **(H)** tumors. Resistant HER2-high tumors display increased communication involving tumor cells and multiple immune cell populations, highlighting broad reorganization of the cellular interaction landscape during therapeutic resistance.

In HER2-low tumors, resistance was characterized primarily by increased interaction diversity with minimal change in overall signaling strength, consistent with rewiring of DC/myeloid-centered crosstalk. In HER2-high tumors, resistance involved increases in both interaction number and signaling strength, indicating broader amplification of tumor–immune communication. These findings suggest that resistance emerges through subtype-specific remodeling of immune function and intercellular signaling, with HER2-low tumors relying more on microenvironmental reorganization and HER2-high tumors showing stronger global communication activation.

## DC trajectory analysis reveals plasticity among homeostatic, IFN-activated, classical cDC2, pDC, and monocyte-derived DC states

Monocle 2 trajectory analysis^60^ resolved the dendritic-cell compartment into a branched pseudotime continuum composed of five transcriptionally defined DC populations (**Figure 4A-D**). The trajectory was organized into seven pseudotime states, indicating that tumor-associated DCs do not exist as isolated subsets but rather occupy interconnected transitional and terminal states. The shared backbone likely represents intermediate DC states, whereas the branch termini reflect divergence toward distinct functional endpoints (**Figure 4A-D**). Subset-specific trajectory mapping showed non-uniform distribution of DC populations across the trajectory. pDCs were concentrated predominantly within a terminal branch and were largely assigned to a single dominant pseudotime state, consistent with a relatively discrete pDC endpoint. In contrast, homeostatic cDC2 cells were broadly distributed across multiple regions of the trajectory, suggesting a more plastic or intermediate homeostatic state. IFN-activated mature cDCs and classical functional cDC2 cells populated intermediate-to-late branches, consistent with acquisition of inflammatory and antigen-presentation programs. ITGAX^+^ monocyte-derived DCs also occupied late trajectory regions, but along a distinct branch, supporting their divergence toward a myeloid-derived activated state. Density analysis (**Figure 4E**) across pseudotime^61^ further highlighted these differences. Homeostatic cDC2 cells showed a broad distribution spanning early to late pseudotime, whereas IFN-activated mature cDCs, classical functional cDC2 cells and ITGAX^+^ monocyte-derived DCs were enriched in intermediate and late phases. pDCs showed the strongest accumulation toward the late end of pseudotime, supporting their localization at a terminal branch. Pie-chart analysis of state composition (**Figure 4F)** was concordant with these distributions, demonstrating that each DC subset contributes preferentially to specific pseudotime states rather than being evenly represented across the entire continuum. Pseudotime-regulated gene analysis by heatmap^61^ identified multiple transcriptional modules associated with progressive DC-state transitions (**Figure 4G**). These modules were enriched for biological processes related to apoptotic signaling, protein kinase activity, ATP binding, immune response, chemotaxis, inflammatory response, chromatin organization, innate immunity, cell adhesion and collagen fibril organization. Collectively, these patterns indicate that DC progression is accompanied by coordinated remodeling of survival, activation, migratory and immune-regulatory programs as shown by GO **(Supplementary Figure 6**) and KEGGs **(Supplementary Figure 7)**. Together, these findings define a dynamic developmental and functional landscape for tumor-associated DCs.

**Figure 4.**
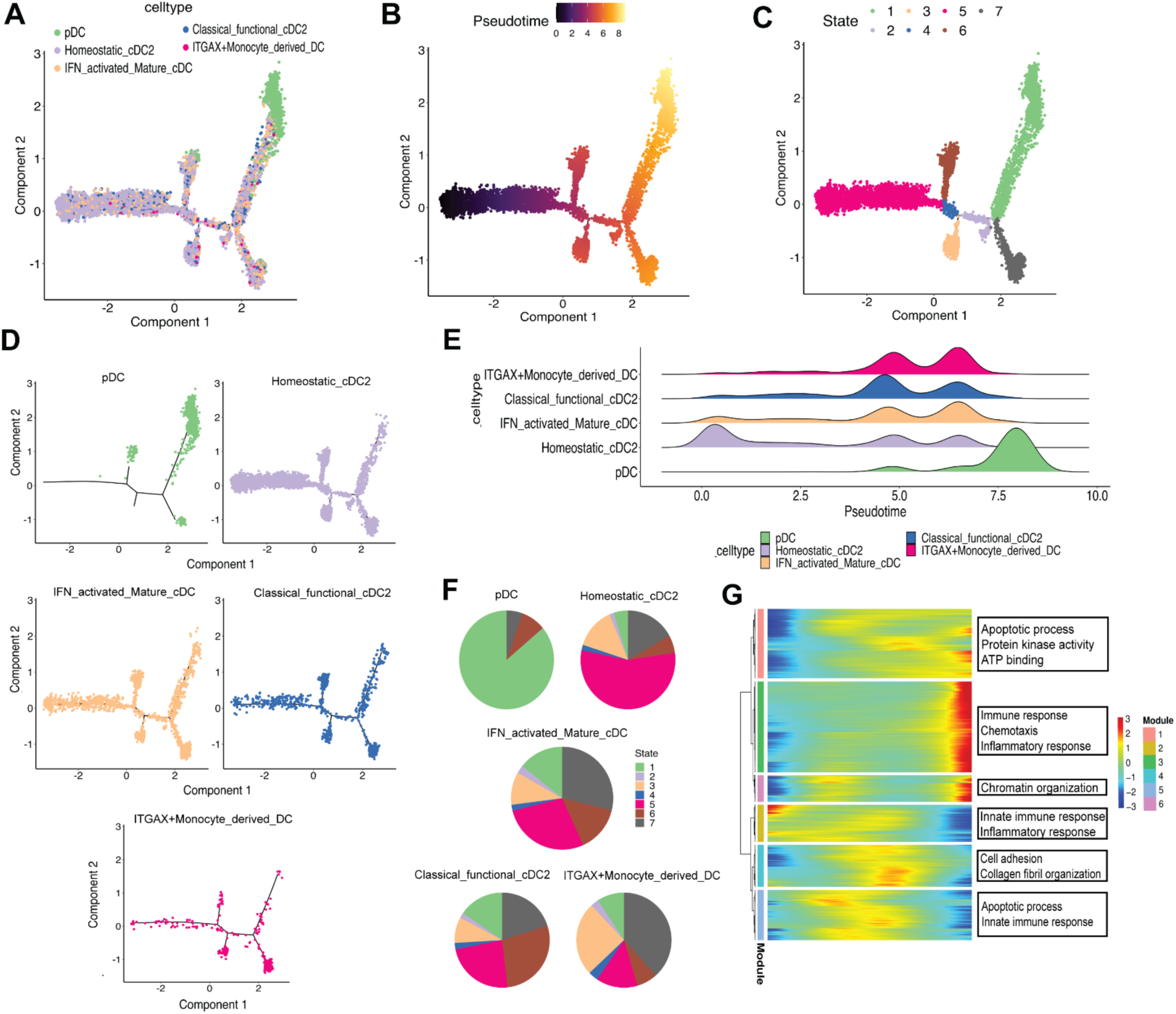
**Trajectory analysis reveals dynamic differentiation and functional transitions among dendritic- cell populations in HER2 breast tumors**. **(A)** Monocle trajectory analysis of dendritic cells colored by cell type. Five transcriptionally distinct dendritic-cell populations—including plasmacytoid dendritic cells (pDCs), homeostatic cDC2, IFN-activated mature cDCs, classical functional cDC2, and ITGAX⁺ monocyte-derived dendritic cells—are distributed along an inferred transcriptional trajectory, indicating progressive state transitions. **(B)** Pseudotime ordering of dendritic cells. Cells are colored according to pseudotime progression, illustrating a continuous differentiation trajectory from early to late cellular states. **(C)** Monocle trajectory colored by inferred cell states. Seven discrete transcriptional states are identified along the developmental continuum, highlighting progressive changes in dendritic-cell identity during differentiation. **(D)** Subset-specific trajectory analysis of individual dendritic-cell populations. Independent trajectory reconstruction demonstrates the developmental positioning and lineage relationships of pDCs, homeostatic cDC2, IFN-activated mature cDCs, classical functional cDC2, and ITGAX⁺ monocyte-derived dendritic cells within the overall pseudotime landscape. **(E)** Density distribution of each dendritic-cell subset across pseudotime. Ridge plots illustrate preferential enrichment of individual DC populations at distinct stages of the developmental trajectory, revealing dynamic temporal changes in cell-state abundance. **(F)** Pie charts show the proportional contribution of pseudotime states within each dendritic-cell population. Distinct DC subsets occupy unique combinations of transcriptional states, demonstrating heterogeneity in differentiation status and functional maturation. **(G)** Heatmap of genes dynamically regulated during pseudotime. Rows represent pseudotime-dependent genes and columns represent cells ordered by pseudotime. Gene expression modules reveal sequential activation and repression of biological programs during dendritic-cell differentiation. Representative Gene Ontology (GO) enrichment analysis identifies pathways associated with immune activation, cytokine-mediated signaling, inflammatory responses, leukocyte migration, antigen processing and presentation, lymphocyte activation, regulation of cell adhesion, and adaptive immune responses.

### Spatial neighborhoods and niches define immune-accessible and resistant architectures

Spatial niche analysis identified 16 tumor microenvironmental states that were differentially enriched across HER2-low and HER2-high tumors according to treatment response (**Figure 5A-C**). HER2-low sensitive tumors **(Figure 5D, E, G**) were enriched for Niche 2, Niche 3, Niche 8, Niche 12 and Niche 16, reflecting a more spatially admixed architecture with tumor, immune and stromal compartments distributed across the tissue. In contrast, HER2-low resistant tumors (**Figure 5D, E, G**) were enriched for Niche 9, Niche 11, Niche 13 and Niche 15, and showed larger, more organized niche territories, consistent with stromal/myeloid-associated remodeling and increased tumor–microenvironment compartmentalization. HER2-high tumors exhibited a distinct spatial organization. Sensitive HER2-high tumors (**Figure 5D, F, H)** were enriched for Niche 1, Niche 5, Niche 9 and Niche 14, whereas resistant HER2-high tumors (**Figure 5D, F, H**) showed broader enrichment of Niche 3, Niche 10, Niche 11, Niche 12, Niche 13 and Niche 15. The overlap of Niche 11, Niche 13 and Niche 15 between HER2-low and HER2-high resistant tumors suggests that shared resistant spatial programs may emerge across HER2 states, while subtype-specific niches indicate distinct routes to therapy escape.

**Figure 5.**
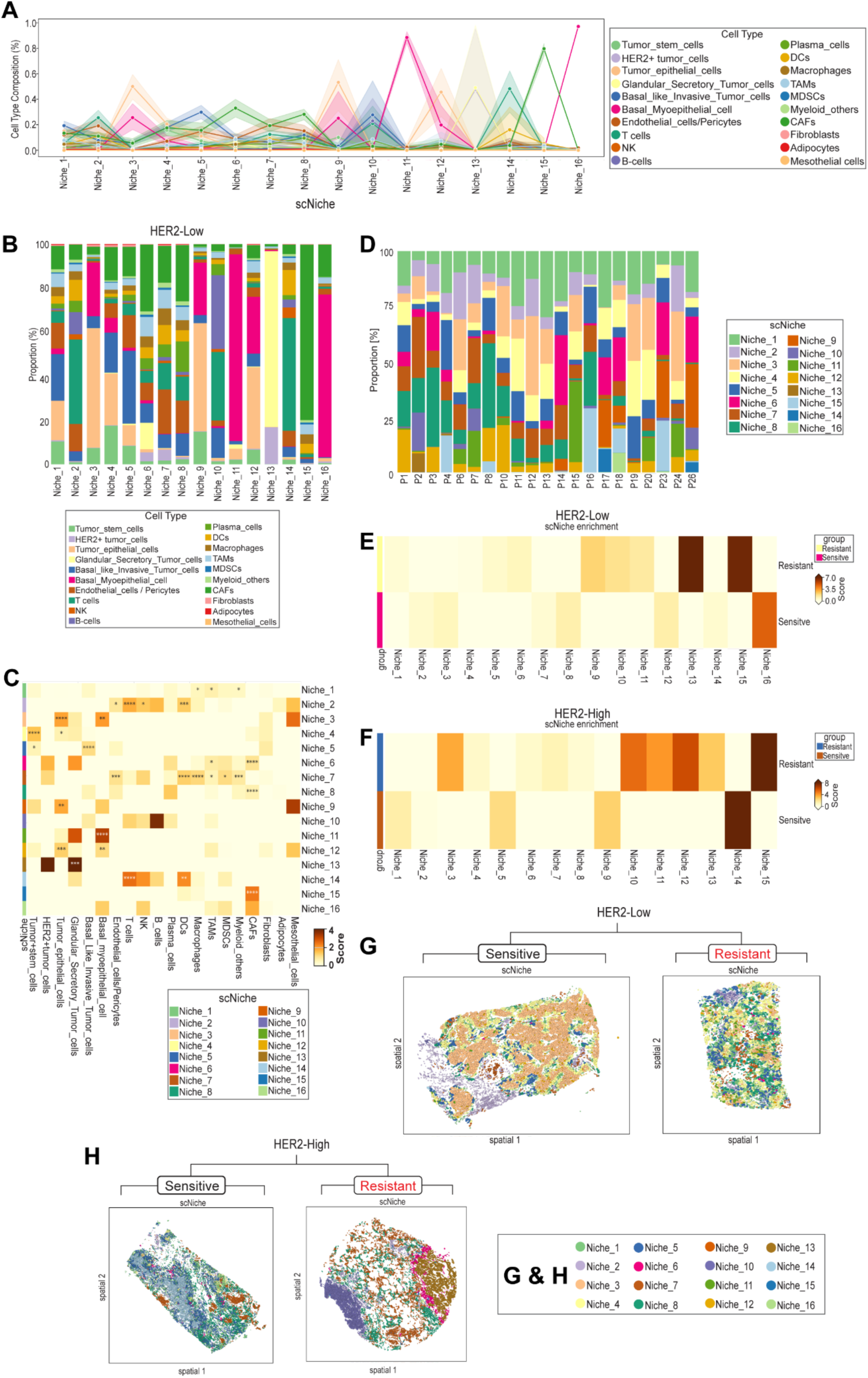
Spatial niche architecture identifies resistance-associated cellular neighborhoods in HER2-low and HER2-high breast tumors. **(A)** Cellular composition of the identified spatial niches. Line plot showing the relative abundance of major tumor, immune, stromal, and vascular cell populations across the spatial niches identified by scNiche analysis, illustrating the distinct cellular composition of each neighborhood. **(B)** Distribution of spatial niches across individual HER2-low tumors. Stacked bar plots represent the relative abundance of each niche in sensitive and resistant patient samples, highlighting inter-patient heterogeneity and subtype-specific niche composition. **(C)** Heatmap showing the enrichment of individual cell populations within each spatial niche. Distinct niches are characterized by unique combinations of tumor, immune, stromal, and vascular cells, demonstrating that each niche represents a specialized multicellular microenvironment. **(D)** Distribution of spatial niches across individual HER2 patients. Stacked bar plots illustrate the relative abundance of each niche in sensitive and resistant tumors, revealing subtype-specific remodeling of spatial neighborhood organization. **(E)** Spatial niche enrichment analysis in HER2-low tumors. Heatmap showing the relative enrichment score of each spatial niche in sensitive and resistant tumors. Sensitive tumors are enriched for a subset of interaction-rich niches, whereas resistant tumors preferentially occupy distinct resistance-associated spatial neighborhoods. **(F)** Spatial niche enrichment analysis in HER2-high tumors. Heatmap illustrating differential enrichment of spatial niches between sensitive and resistant tumors. Resistant HER2-high tumors exhibit selective expansion of specific spatial neighborhoods compared with sensitive tumors. **(G)** Representative spatial maps of HER2-low tumors showing the tissue distribution of scNiche-defined cellular neighborhoods in sensitive and resistant tumors. Sensitive tumors display heterogeneous, spatially distributed niches, whereas resistant tumors exhibit expansion and consolidation of resistance-associated niches. **(H)** Representative spatial maps of HER2-high tumors illustrating niche organization in sensitive and resistant tumors. Resistant tumors demonstrate remodeling of spatial neighborhood architecture, with preferential expansion of specific cellular niches compared with sensitive tumors.

Together, these data demonstrate that therapeutic response is associated with spatial organization of the tumor ecosystem. HER2-low resistance is marked by stromal/myeloid-enriched and compartmentalized niches, whereas HER2-high resistance is associated with broader spatial diversification and epithelial–stromal remodeling. These findings support scNiche architecture as a spatial biomarker framework for identifying therapy-sensitive and therapy-resistant tumor states.

## Spatial imaging reveals distinct immune microenvironmental states in HER2-low sensitive and resistant tumors

To further define how treatment response is associated with spatial organization of the tumor microenvironment, we analyzed the cellular landscape and in situ gene expression analysis with Xenium (**Figure 6A).** This analysis revealed marked differences in immune-cell infiltration, stromal organization, and tumor-cell architecture between sensitive and resistant tumors. HER2-low sensitive tumors displayed an immune-inflamed phenotype, with CD8 T cells, DCs, NK cells and TAMs interspersed within tumor epithelial regions, consistent with preserved tumor–immune contact. In contrast, HER2-low resistant tumors exhibited immune-excluded or immune-desert architectures (**Figure 6B**), characterized by CAF-rich stromal organization, basal-like tumor-cell enrichment and reduced CD8 T-cell infiltration. The tumor microenvironmental organization of HER2-low sensitive and resistant tumors is shown in **Figure 6C**, including spatial distribution of immune cells marked by CD8, CAFs marked by FAP^62^, and tumor epithelial cells marked by CDH1. Spatial organization of the tumor microenvironment further revealed marked differences between therapy-sensitive and therapy-resistant tumors. Sensitive tumors displayed a more immune-infiltrated architecture, characterized by greater abundance and intratumoral distribution of CD8A-positive cytotoxic T cells, consistent with preserved immune accessibility and active antitumor surveillance (**Figure 6D).** In contrast, resistant tumors showed reduced CD8A-positive immune infiltration and a more immunosuppressive, invasive, and stromally remodeled microenvironment. These tumors exhibited increased expression of CD44, indicating enrichment of cancer stem-like cells; CD274, reflecting elevated PD-L1-mediated immune evasion; MMP9, consistent with enhanced invasive potential; and FAP, marking expansion of the CAF compartment **Figure 6D)**. Marker-based spatial profiling further identified heterogeneous DC states (**Figure 6E),** including pDC, homeostatic cDC2, IFN-activated mature cDC, classical functional cDC2 and ITGAX^+^ monocyte-derived DC populations, distributed alongside tumor epithelial cells, CAFs, T cells, NK cells and endothelial cells (**Figure 6F**). Cytokine mapping showed spatially localized IL10, IL12A, IL12B and TNF signals, indicating co-existing inflammatory and immunoregulatory programs (**Figure 6G**). Together, these findings suggest that therapy sensitivity is associated with immune-accessible spatial organization, whereas resistance is linked to CAF-rich immune exclusion, immune-cold remodeling and altered DC/myeloid organization. Overall, these data indicate that HER2-low resistant tumors are not defined simply by a reduction in immune cells, but by a broader spatial reorganization of the tumor microenvironment. Resistant tumors show increased CAF accumulation, reduced CD8 T-cell infiltration, enrichment of suppressive myeloid populations (TAMs), and expansion of basal-like invasive tumor-cells. These findings support the concept that spatial immune exclusion and stromal remodeling are key features of therapeutic resistance in HER2-low breast cancer. Multiplex immunostaining **(Figure 6H)** revealed that sensitive tumors were enriched in CD8A⁺ T cells, NK cells, and M1 macrophages, whereas resistant tumors exhibited markedly reduced infiltration of these antitumor immune populations, consistent with an immune-suppressed and immune-excluded tumor microenvironment.

**Figure 6.**
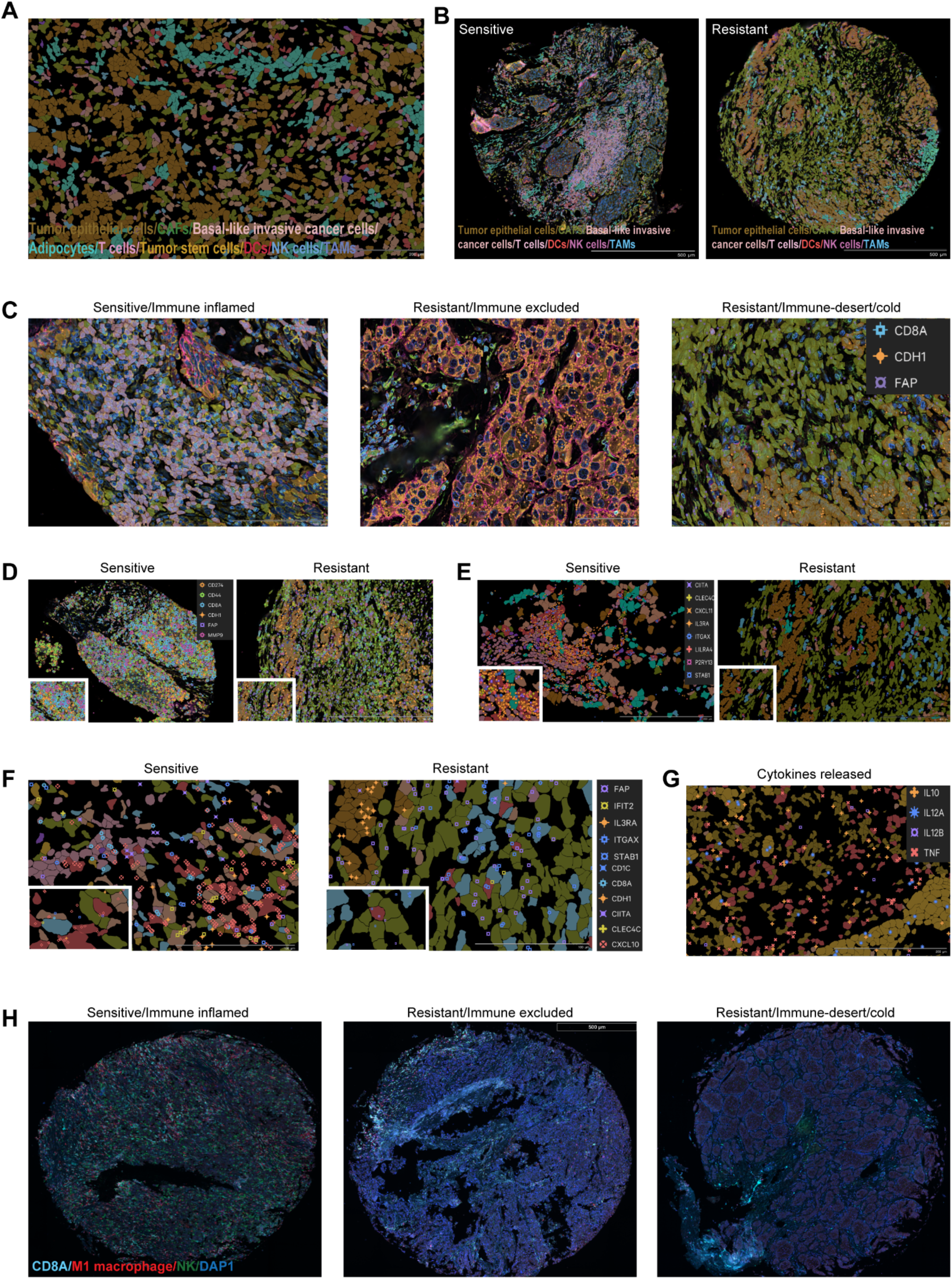
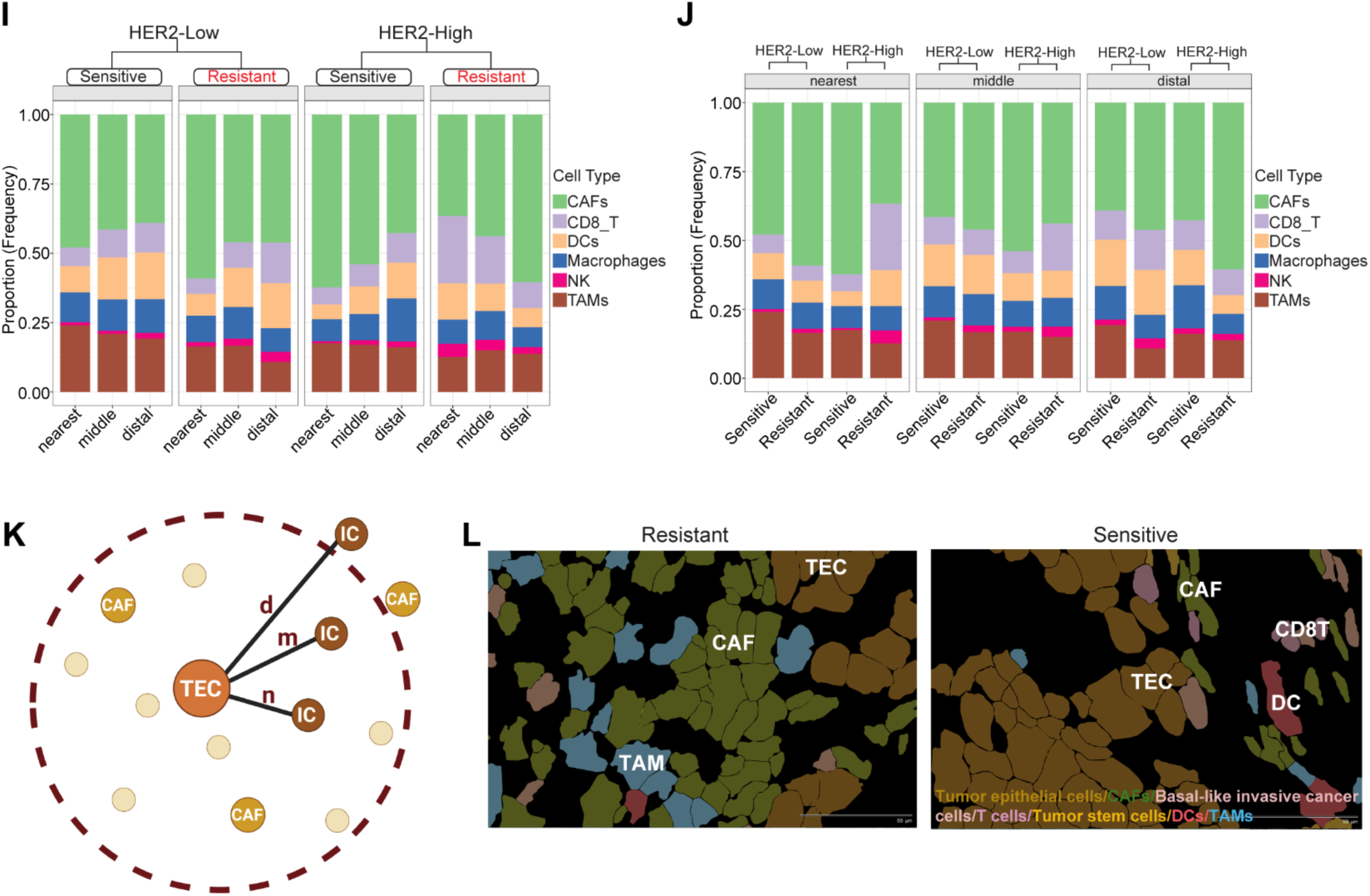
Spatial organization of immune and tumor cell populations defines distinct immune landscapes associated with therapeutic response. **(A)** Representative multiplex spatial transcriptomic image illustrating the overall organization of major cellular populations within the HER2-low tumor microenvironment, including tumor epithelial cells, basal-like invasive tumor cells, CAFs, DCs, NK cells, T cells, tumor stem cells, adipocytes, and TAMs. **(B)** Representative whole-tissue spatial maps of HER2-low sensitive and resistant tumors demonstrating global differences in tissue architecture and cellular organization. Resistant tumors exhibit increased spatial compartmentalization and reduced immune infiltration compared with sensitive tumors. **(C)** Representative high-resolution images illustrating three distinct immune phenotypes identified in HER2-low tumors: immune-inflamed (sensitive), immune-excluded (resistant), and immune-desert (cold) (resistant). **(D)** Spatial distribution of major cellular populations with the selective markers in representative HER2-low sensitive and resistant tumors. Sensitive tumors exhibit heterogeneous intermixing of tumor and immune cells, whereas resistant tumors demonstrate increased spatial segregation and enrichment of stromal and tumor compartments. Insets highlight representative cellular neighborhoods. **(E)** Spatial localization of dendritic-cell populations in HER2-low tumors using subtype-specific marker genes. **(F)** Spatial localization of different dendritic cell population along with the tumor immune microenvironment. **(G)** Spatial expression of representative cytokines within HER2-low tumors, demonstrating localized inflammatory signaling within distinct tissue regions. Cytokine expression patterns highlight regional heterogeneity in immune activity across the tumor microenvironment. **(H)** Multiplex immunofluorescence validation of immune phenotypes using CD8A, M1 macrophage, NK-cell, and DAPI staining. Sensitive tumors exhibit an immune-inflamed phenotype characterized by abundant CD8⁺ T cells, M1 macrophages, and NK cells throughout the tumor. In contrast, resistant tumors display either immune-excluded architecture, with immune cells confined to the tumor periphery, or immune-desert architecture characterized by markedly reduced immune-cell infiltration. **(I-J)** Relative composition of CAFs, CD8⁺ T cells, DCs, macrophages, NK cells, and TAMs within nearest (0–20 μm), middle (20–40 μm), and distal (>40 μm) spatial neighborhoods surrounding tumor epithelial cells in HER2-low and HER2-high tumors. Stacked bar plots illustrate changes in the spatial distribution of immune and stromal populations across tumor-proximal and tumor-distal regions. Schematic representation of spatial proximity **(K)** and the distance distribution of individual cell populations relative to tumor epithelial cells. **(L)** Spatial images show the presence of different immune cells at increasing distances from tumor epithelial cells. CAFs and macrophages are preferentially localized near tumor cells, whereas CD8⁺ T cells, dendritic cells, and NK cells exhibit broader spatial distributions. TEC: Tumor epithelial cells, IC: Immune cells, CAF: Cancer-associated fibroblasts, N: Nearest, M: Middle, D: Distal.

Finally, our spatial proximity analysis **(Figure 6I-L)** demonstrates that immune-cell abundance alone is insufficient to explain therapeutic response. Instead, the relative positioning of immune cells with respect to tumor epithelial cells represents a critical determinant of treatment outcome. HER2-low sensitive tumors maintain close spatial associations between tumor epithelial cells (**Figure 6I-L**) and key immune populations, whereas resistant tumors display increased immune-cell displacement into more distal regions. Similar trends are observed in HER2-high tumors, although the degree of spatial remodeling appears less extensive **(Figure 6I-L)**. These findings suggest that spatial accessibility of immune cells to tumor cells is an important architectural feature of chemotherapy response and provide quantitative evidence that therapeutic resistance is accompanied by reorganization of the tumor immune landscape.

Spatial signature analysis revealed distinct microenvironmental organization between HER2-low sensitive and resistant tumors. Sensitive tumors were characterized by a more admixed spatial architecture, with immune and tumor compartments distributed across the tissue, suggesting greater immune accessibility and preserved tumor–immune interaction. In contrast, resistant tumors showed a more compartmentalized spatial pattern, with enrichment of stromal/myeloid-associated niches and stronger tumor–microenvironment segregation (**Figure 7).**

**Figure 7.**
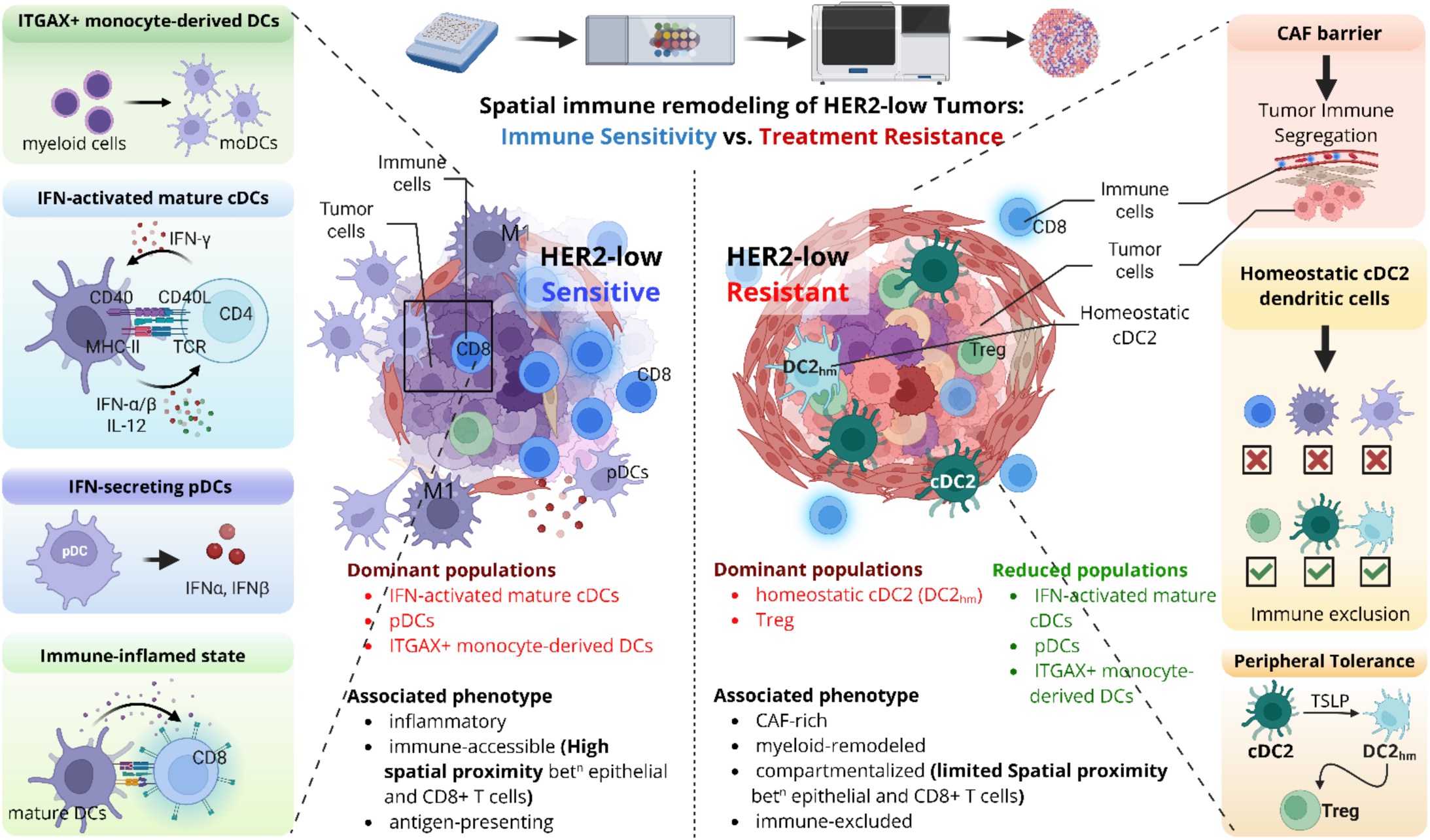
Spatial signature underlying chemotherapy response in HER2-low breast cancer. Schematic overview illustrating the spatial and functional organization of the tumor microenvironment associated with chemotherapy sensitivity and resistance in HER2-low breast tumors.

## Discussion

HER2-low breast cancer is biologically heterogeneous, and the determinants of therapeutic response extend beyond tumor-cell HER2 expression. Using spatial transcriptomic profiling of clinically annotated HER2-low and HER2-high breast tumors, this study shows that treatment resistance is associated with coordinated remodeling of dendritic-cell states, tumor–immune communication, and spatial niche organization. Sensitive tumors displayed a more immune-accessible architecture, whereas resistant tumors were characterized by CAF-rich stroma, myeloid remodeling, reduced CD8 T-cell access, and greater compartmentalization of tumor and immune regions.

Dendritic cells are important regulators of the breast cancer TME because they control how tumor antigens are presented to T cells and whether the immune system mounts an effective antitumor response. In therapy-sensitive tumors, DC can support immune activation by promoting antigen presentation, interferon signaling, chemokine production, and recruitment of CD8 T cells and NK cells. This can create an immune-inflamed tumor microenvironment that improves response to chemotherapy or other therapies. However, in drug-resistant tumors, DCs may become functionally remodeled. Instead of supporting productive antitumor immunity, resistant tumors may show enrichment of DC states associated with chronic inflammation, immune suppression, hypoxia, lactate metabolism, wound healing, and myeloid remodeling. These altered DC states may fail to effectively activate cytotoxic T cells and may instead contribute to an immune-suppressed or therapy-protective tumor niche. A major finding was the functional heterogeneity of the dendritic-cell compartment. Five distinct DC states were identified, including classical functional cDC2, homeostatic cDC2, IFN-activated mature cDC, ITGAX^+^ monocyte-derived DC, and pDC populations. Resistant HER2-low tumors showed increased homeostatic and classical cDC2 states, together with reduced IFN-activated mature DCs and pDCs. Importantly, independent validation in the TCGA-BRCA cohort demonstrated that homeostatic cDC2 signatures were associated with poor survival. These data suggest that resistant tumors retain antigen-presenting DC populations but may fail to sustain the interferon-driven inflammatory programs required for effective CD8 T-cell and NK-cell activation^63^. In contrast, sensitive tumors were enriched for pDC and ITGAX^+^ monocyte-derived DC states, potentially reflecting a more inflammatory and therapy-responsive immune environment^39^.

Homeostatic DCs likely represent a tissue-surveillance dendritic-cell state^40^. They are not fully inflammatory or mature, but they retain important functions such as antigen uptake, receptor-mediated endocytosis, MHC-II antigen processing/presentation, and communication with immune and stromal cells^64^. In tumors, this state can have a dual role: it may support immune monitoring in sensitive tumors, but it may also become a reservoir that is redirected into suppressive or ineffective DC states during therapy resistance^65,66^. In HER2-low tumors, homeostatic cDC2 forms the major DC backbone, meaning it may be the starting or transitional DC state from which other DC programs emerge. In sensitive tumors, homeostatic cDC2 may transition toward IFN-activated mature cDC and classical functional cDC2 states, supporting antigen presentation, chemokine signaling, CD8 T-cell recruitment, and productive antitumor immunity. This aligns with the known role of conventional DCs in antigen presentation and T-cell activation, including cDC2 involvement in CD4 T-cell activation and durable cytotoxic T-cell responses^13,50^. In resistant tumors, however, homeostatic DCs may be functionally diverted. Instead of maturing into immunostimulatory DC states, they may shift toward pDC-like or ITGAX^+^ monocyte-derived DC-like programs associated with hypoxia, lactate metabolism, wound healing, TLR/MAPK signaling, and dysfunctional tumor–immune communication. Tumor-infiltrating DCs are known to be plastic and can switch from immunostimulatory to inhibitory/tolerogenic roles during cancer progression^65^.

Trajectory analysis further indicated that these DC populations exist along a dynamic continuum rather than as fixed subsets. Homeostatic cDC2 cells occupied broad transitional regions, whereas IFN-activated mature DCs, classical cDC2 cells, monocyte-derived DCs, and pDCs localized to more differentiated branches. This suggests that treatment response may depend not only on the abundance of individual DC subsets, but also on the ability of the DC compartment to transition toward productive inflammatory and antigen-presenting states. CellChat analysis showed that HER2-low resistance was associated with a substantial increase in the number of inferred cellular interactions without a major increase in overall signaling strength. Thus, resistant tumors were not communication-deficient; instead, their signaling networks appeared diversified and rewired toward DC-, macrophage-, and myeloid-centered crosstalk. This contrasts with HER2-high resistant tumors, in which both interaction number and strength increased, suggesting that resistance mechanisms differ across HER2-defined disease states.

Spatial niche analysis showed that, sensitive HER2-low tumors showed a more intermixed distribution of tumor, immune, and stromal cells, potentially facilitating antigen presentation and direct immune-cell access to malignant regions. Resistant tumors contained larger and more organized stromal/myeloid niches, consistent with the formation of protected tumor territories. The enrichment of CAFs, TAMs, basal-like invasive tumor cells, and reduced CD8 infiltration supports a model in which stromal remodeling and myeloid suppression cooperate to limit effective antitumor immunity. Importantly, resistant tumors were not uniformly immune-deserted. Instead, inflammatory and immunoregulatory signals coexisted within spatially distinct regions. This suggests that resistance may arise from dysfunctional or mislocalized immune activation rather than complete immune absence. The presence of DCs and other immune cells in resistant tumors therefore does not necessarily indicate effective immunity; their activation state and spatial relationship with tumor cells are likely to be more informative than abundance alone. Together, these findings support a model in which HER2-low resistance emerges through spatial reorganization of the tumor ecosystem. Reduced IFN-activated DC programs, increased homeostatic and classical cDC2 states, expanded myeloid communication, CAF-associated immune exclusion, and enrichment of aggressive tumor-cell states collectively create a microenvironment that permits residual tumor survival after therapy. These results suggest that spatial measurements of DC-state composition, tumor–immune proximity, and resistant niche abundance may provide useful biomarkers of treatment response. The distance between tumor epithelial cells and key immune populations—including dendritic cells and CD8⁺ T cells—reflects the functional capacity of the tumor microenvironment to support anti-tumor immunity. Chemotherapy-sensitive tumors maintain close immune–tumor proximity, whereas resistant tumors exhibit increased spatial separation, reducing immune surveillance and contributing to therapeutic resistance.

This study is limited by its retrospective design, cohort heterogeneity, and reliance on inferred communication and transcriptionally defined cell states. Larger independent cohorts and orthogonal validation by spatial proteomics, and functional assays will be required. Nevertheless, the data establish that HER2-low therapeutic resistance is associated with coordinated dendritic-cell, stromal, and spatial remodeling rather than with a single dominant cellular mechanism. In summary, sensitive HER2-low tumors retain an immune-accessible and inflammatory architecture, whereas resistant tumors develop a CAF-rich, myeloid-organized, and spatially compartmentalized microenvironment. These findings identify dendritic-cell state and immune–stromal spatial organization as potential biomarkers and therapeutic vulnerabilities in HER2-low breast cancer.

## Methods

### Sample collection

Patients were retrospectively identified under MD Anderson protocol 2024-0158, which includes consent for the collection of archival tissue samples. Eligible cases were selected based on the availability of tissue specimens within our institutional tissue bank. Formalin-fixed paraffin-embedded (FFPE) tumor tissue blocks were obtained under an MD Anderson Cancer Center institutional review board–approved protocol (2021-0764), with informed patient consent confirmed before use. Corresponding hematoxylin and eosin (H&E)–stained sections were scanned (Aperio AT2 and GT450, Leica Biosystems) and reviewed using digital pathology workflows. Whole-slide images were examined using ImageScope (Leica Biosystems) and HALO (Indica Labs) to verify tumor presence, assess spatial distribution, and identify regions enriched for residual tumor. Tumor-containing areas were annotated on H&E sections, and blocks were evaluated for suitability for downstream spatial transcriptomic analysis. Tissue microarrays (TMAs) were generated from FFPE specimens from 21 patients. TMAs were constructed using the Grand Master automated tissue arrayer (Epredia), incorporating 1-mm cores into two recipient blocks (80 cores total). Completed TMA blocks and corresponding slides underwent quality control to confirm tissue integrity and adequate tumor representation. Sections (5 µm) were cut from each TMA block and mounted onto Xenium slides following the manufacturer’s instructions. Slides were incubated, dried, and stored in a desiccator at room temperature until processing. H&E-stained TMA sections were digitally scanned and annotated by a research pathologist using QuPath^67^. Annotated images were imported into Xenium Explorer for registration and alignment with spatial transcriptomic data. Probe hybridization, signal amplification, and library preparation were performed using the Xenium Spatial Gene Expression Reagent Kits for FFPE (10x Genomics) according to the manufacturer’s protocols. Sequencing was performed on a NovaSeq 6000 platform (Illumina).

### Digital pathology

Selected tissue sections were digitized using an Aperio GT450 (Leica Biosystems, Illinois, USA) whole-slide scanner. Digital whole-slide images were reviewed for image quality, tissue completeness, focus, and suitability for analysis. Scanned slides were used for digital pathology assessment, including confirmation of tumor regions and evaluation of tissue adequacy for downstream image-based analysis. Regions containing viable tumor were prioritized, while areas with extensive necrosis, poor fixation, tissue folds, scanning artifacts, or insufficient tumor cellularity were excluded when applicable. Digital images were reviewed in conjunction with the corresponding histologic sections to ensure accurate tumor annotation and consistency across cases. The final digital pathology dataset included only slides that passed quality control and contained adequate representative tumor tissue for downstream assays and analysis.

### Xenium in situ workflow

Human formalin-fixed paraffin-embedded (FFPE) tissue sections were profiled using the Xenium Prime 5K Gene Expression assay with cell segmentation on the Xenium Analyzer (10x Genomics, Pleasanton, CA, USA). Samples were processed using Xenium slides, Xenium Prime Sample Prep Reagents (PN-1000720), Xenium Prime Cassettes and Inserts (PN-1000723), the Xenium Prime 5K Human Pan Tissue and Pathways Panel (PN-1000724), Xenium Cell Segmentation Staining Reagents (PN-1000661), and Xenium Prime 5K decoding reagents and consumables according to the manufacturer’s instructions, unless otherwise noted. No custom add-on panel was used.

### Xenium sample preparation

FFPE blocks were sectioned at 5 µm and mounted within the sample area of Xenium slides. Slides were equilibrated to room temperature for 30 min, air-dried overnight at room temperature, baked at 42 °C for 3 h, and stored in a desiccated environment until processing. Immediately before deparaffinization, slides were baked at 60 °C for 120 min. Slides were then deparaffinized by sequential immersion in xylene (2 × 10 min), 100% ethanol (2 × 3 min), 95% ethanol (2 × 3 min), 70% ethanol (1 × 3 min), and nuclease-free water (20 s). After rehydration, slides were immediately assembled into Xenium Prime cassettes to prevent tissue drying. Tissue sections were decrosslinked using Xenium Prime Sample Prep Reagents according to the manufacturer’s protocol, including incubation at 80 °C for 30 min. Sections then underwent priming hybridization at 50 °C for 90 min, followed by a post-priming wash at 50 °C for 30 min, RNase treatment at 37 °C for 20 min, and polishing at 37 °C for 1 h. Probe hybridization with the Xenium Prime 5K Human Pan Tissue and Pathways Panel was performed at 50 °C for 16 h.

On the following day, sections were subjected to a post-hybridization wash at 35 °C for 15 min, ligation at 42 °C for 30 min, amplification enhancement at 4 °C for 2 h, and amplification at 30 °C for 90 min. For cell segmentation, amplified sections were processed using Xenium Cell Segmentation Staining Reagents. Sections were equilibrated through 70% ethanol, 100% ethanol, 100% ethanol, 70% ethanol, and PBS-T, blocked in diluted Xenium Block and Stain Buffer for 1 h at room temperature, and incubated with Xenium Multi-Tissue Stain Mix overnight at 4 °C. On the following day, sections underwent stain enhancement for 20 min at room temperature, autofluorescence quenching with diluted Reducing Agent B for 10 min followed by AF Solution for 10 min, drying at 37 °C for 5 min, and nuclei staining for 1 min at room temperature. Slides were then loaded onto the Xenium Analyzer with the appropriate decoding reagents and consumables, and imaging, transcript decoding, and cell segmentation were performed according to the manufacturer’s instructions.

### Post xenium H& E staining

After Xenium imaging, slides were removed from the Xenium cassette and processed for hematoxylin and eosin (H&E) staining. Residual quencher was removed by incubating slides for 10 min at room temperature in freshly prepared quencher removal solution consisting of 69.6 mg sodium hydrosulfite dissolved in 40 ml molecular-grade water, followed by three 1-min washes in molecular-grade water. Slides were either processed immediately for H&E staining or stored in 1× PBS at 4 °C for up to 2 days before staining. For H&E staining, slides were immersed in molecular-grade water for 2 min, stained in Mayer’s hematoxylin for 20 min, washed in molecular-grade water three times for 1 min each, differentiated in Dako Bluing Solution for 1 min, and rinsed in molecular-grade water for 1 min. Sections were then dehydrated through 70% ethanol for 3 min and 95% ethanol for 3 min, stained in Eosin Y, alcoholic for 4.5 min, and further dehydrated through 95% ethanol (2 × 30 s) and 100% ethanol (2 × 30 s). Slides were cleared in xylene (2 × 3 min), coverslipped, and dried in a fume hood for 30 min. H&E-stained slides were scanned at 20× magnification using a Leica Aperio CS2 whole-slide scanner.

### Multiplexing immunostaining on TMAs

Antibodies: Two separate panels were developed and optimized for use in this study. The following primary antibodies were utilized for both immunohistochemical and immunofluorescence staining: rabbit anti-human CD56 [BLR152J], rabbit anti-human CD86 [E2G8P], and mouse anti-human CD8a [144B]. Optimization: Optimization of antibody concentration and staining order was achieved using FFPE human immune tissue microarray serial sections. Each target was evaluated for overall signal: noise ratio, loss of signal intensity, elution efficiency, and overall autofluorescence via heat-induced epitope retrieval (HIER) methods. The final optimized panels were then subjected to the Spatomics’ CFP™ method of multiplex immunofluorescence (see below). Multiplex immunofluorescence staining was performed with Spatomics CFP™ cleavable dyes. This technology allows for numerous protein targets to be stained using CFPs, which generate strong, localized fluorescence through an HRP-catalyzed reaction between the dye and nearby residues. After imaging, the fluorescent tags are efficiently cleaved and HRP is deactivated, allowing repeated cycles without compromising tissue quality or antigenicity. FFPE microarrays were baked for 60 minutes, deparaffinized in xylene, and rehydrated by serial passage through graded concentrations of ethanol. Endogenous peroxidase in tissues was blocked with 0.9% H2O2/methanol for 40 min. An initial HIER treatment was performed for 20 min at 92-96C in Tris EDTA pH9 buffer. Following HIER, slides were rinsed with DI water and cooled at RT for 20 minutes. Slides were blocked with 20% normal goat serum for 30 minutes before loading onto the Parhelia Spatial Station™ auto stainer. Primary antibodies were incubated for 20 min. Then, slides were rinsed with TBS for 10 min and incubated with HRP-conjugated secondary (A120-501P) for 20 min, followed by another 10 min rinse in TBS. Incubation with a CFP dye was done for two 10 min exchanges, followed by a 10 min rinse in DI water. After the first panel was stained, the slide was removed and incubated with DAPI for 10 minutes. The slide was mounted in VECTASHIELD Vibrance Antifade Mounting Medium (ThermoFisher Scientific) and imaged using the PhenoImager HT. Following imaging, the coverslip was removed and the first panel of bound primary and secondary antibodies was then cleaved off the tissue. The staining and imaging process was repeated for the second panel, resulting in two images [Panel 1 and Panel 2] of the same slide. Imaging: Akoya Biosciences’ PhenoImager HT (formerly known as Vectra Polaris Automated Quantitative Pathology Imaging System) was used for multispectral imaging at 40× magnification. Thereafter, whole slide images were uploaded for viewing on Pathcore.

### Xenium In Situ Gene Expression Data Processing

Xenium Onboard Analysis (version 4.0.1.0) pipeline supports the steps of transcript decoding and cell segmentation; a quality score is generated for each transcript. Only high-quality transcripts with Q-Score ≥ 20 were retained to construct the cell-feature matrix. The cell-feature matrix was then analyzed using the Scanpy (version 1.11.1) Python package. To remove low-quality cells, the following criteria were applied: cells were filtered by (1) gene numbers (gene numbers < 5) and (2) transcript numbers (transcript numbers < 10). To obtain the normalized gene expression data, library size normalization was performed using the sc.pp.normalize_total and sc.pp.log1p functions. To integrate spatial information, the BANKSY matrix was generated based on the gene expression and spatial coordinates using Banksy_py (version 1.2.1). Principal component analysis (PCA) was performed to reduce the dimensionality with the sc.pp.pca function. Then, the sc.external.pp.harmony_integrate function was performed to remove batch effects. Graph-based clustering was performed to cluster cells according to the BANKSY matrix with the sc.pp.neighbors and sc.tl.leiden functions. Cells were visualized using a two-dimensional Uniform Manifold Approximation and Projection (UMAP) algorithm with the sc.tl.umap function. The sc.tl.rank_genes_groups (method = "wilcoxon") function was used to identify marker genes of each cluster. Differentially expressed genes (DEGs) were selected using the function sc.tl.rank_genes_groups (method = "wilcoxon"). P value < 0.05 and |log2 fold change| > 1.5 were set as the thresholds for significantly differential expression.

### Cell segmentation

Cell segmentation was performed using the 10x Genomics Xenium Onboard Analysis pipeline with the Xenium Cell Segmentation Staining workflow^68^. Tissue sections were stained with the Xenium Multi-Tissue Stain Mix, and nuclei were identified from DAPI morphology images. Cell boundaries were then inferred using the Xenium multimodal cell segmentation algorithm, which applies deep-learning models to multi- channel morphology stain images. For each cell, segmentation was assigned using a prioritized approach based on boundary stain signal, expansion from the nucleus to the 18S rRNA interior stain edge, or 5 µm nuclear expansion when boundary and interior stain information was insufficient. The resulting cell segmentation masks were used to assign decoded transcripts to cells and generate the cell-feature matrix for downstream analysis.

### Functional and pathway enrichment analysis (GO and KEGG)

The enrichment analysis was performed using an in-house script based on the hypergeometric distribution. All protein-coding genes were used as the background list, and the differentially expressed protein-coding genes were treated as the candidate list. P-values were calculated and then corrected using the Benjamini-Hochberg method. Gene Ontology (GO) enrichment and KEGG pathway enrichment analyses of DEGs were performed using R (version 4.0.3) based on the hypergeometric distribution.

### Monocle2 Pseudotime Analysis

The developmental pseudotime was determined with the Monocle2 package (version 2.9.0)^69^. The raw count data were first converted from a Seurat object into a CellDataSet object using the importCDS function in Monocle. The differentialGeneTest function of the Monocle2 package was used to select ordering genes (qval < 0.01) that were likely to be informative in ordering cells along the pseudotime trajectory. Dimensionality reduction and clustering analysis were performed with the reduceDimension function, followed by trajectory inference with the orderCells function using default parameters. Gene expression was plotted with the plot_genes_in_pseudotime function to track changes over pseudotime.

### Cell Communication Analysis by CellChat

Cell communication analysis was performed using the CellChat (version 2.1.2) R package^70^. First, the normalized expression matrix was imported to create the CellChat object with the createCellChat function. The data were then preprocessed with the identifyOverExpressedGenes, identifyOverExpressedInteractions, and projectData functions using default parameters. The computeCommunProb, filterCommunication (min.cells = 10), and computeCommunProbPathway functions were used to determine potential ligand-receptor interactions. Finally, the cell communication network was aggregated using the aggregateNet function.

### scNiche Analysis

The Python package scNiche (version 1.0.0) was used to identify and characterize cellular niches. First, multi-view features for each cell (cell molecular profile, neighborhood molecular profile, and neighborhood cell composition) were extracted using the process_multi_slices function. The multi-view graph structure was then constructed using the prepare_data function. A multi-view graph neural network model was trained using the Runner class to obtain cell niche embeddings. Finally, clustering was performed using the Leiden clustering method to obtain the niche results.

### TCGA-BRCA Survival Analysis

To investigate whether DC-associated transcriptional programs identified in the spatial analysis were related to clinical outcomes in an independent, broader, publicly available breast cancer cohort, we analyzed RNA-sequencing data from the TCGA-BRCA UCSC Xena HiSeqV2 dataset together with overall-survival information. Five DC marker signatures were evaluated: homeostatic cDC2 (STAB1, IL13RA1, SLCO2B1), plasmacytoid DC (pDC) (CLEC4C, LILRA4, IL3RA), IFN-activated mature cDC (CXCL10, CXCL11, IFIT2), ITGAX^+^ monocyte-derived DC (ITGAX), and classical functional cDC2 (P2RY13, CIITA). Expression of each marker gene was standardized across the cohort to obtain gene-wise z-scores. For multigene signatures, the corresponding z-scores were averaged so that each marker contributed equally to the signature. Samples were then divided into high- and low-score groups using the cohort-specific median signature score. Survival was compared using Kaplan–Meier curves and log-rank tests, with Benjamini–Hochberg correction across the five signatures. A total of 1,214 samples with complete expression and survival information were included.

## Data availability

The Xenium data generated in this study will be made publicly available upon acceptance of the manuscript.

## Code availability

The code will be available upon acceptance of this paper.

## Supporting information

Supplementary doc

## Acknowledgements

This work was funded by the Texas A&M University Faculty Development Grant (to T.R.S.), PRISE grant (to T.R.S.), and Strategic Transformative Research Funds (STRP to T.R.S.), and 1R01GM163238 (to B.M. and T.R.S.). This work was supported, in part, by a Pilot grant from NIEHS P30ES029067 (to T.R.S.). We would like to thank Omics Empower Inc. for helping with the bioinformatics analyses.

## Author contributions

TRS conceived the study; T.R.S., O.O, S.B., S.A., M.S. prepared and organized the data, B.L., Y.X., T.T., B.M. performed bioinformatics data analyses, A.R.S., G.R., D.T., performed pathology data analyses. T.R.S., O.O., S.B., S.A., A.R.S. prepared the figures, legends, and wrote the manuscript. R.S., A.G., and S.T. wrote the manuscript. All authors reviewed, read, and agreed to publish the manuscript.

