## Supplementary doc for "Spatial immune ecosystems govern therapeutic response in HER2-low breast cancer"

**Supplementary Information**

| Patient ID | Age | Race | HER2 Status | ER/PR Status | Neoadjuvant Therapy | Response to treatment |
|---|---|---|---|---|---|---|
| 1 | 48 years | Black or African American | 3+ | <1 | TCHP | Non-pCR |
| 2 | 52 years | Black or African American | 3+ | <1 | TCHP | Non-pCR |
| 3 | 41 years | Black or African American | 3+ | <1 | TCHP | Non-pCR |
| 4 | 43 years | White or Caucasian | 3+ | <1 | TCHP-AC | pCR |
| 6 | 59 years | White or Caucasian | 1+ | <1 | DD-AC-Paclitaxel | pCR |
| 7 | 36 years | Asian | 1+ | <1 | KN-522(pembro-paclitaxel) | Non-pCR |
| 8 | 65 years | White or Caucasian | 1+ | <1 | Sacituzumab+pembro | pCR |
| 10 | 43 years | White or Caucasian | 1+ | 0 | KN-522(pembro-paclitaxel) | pCR |
| 11 | 62 years | White or Caucasian | 2+ | <1 | TCHP | Non-pCR |
| 12 | 68 years | Asian | 2+ | <1 | TCHP | pCR |
| 13 | 55 years | White or Caucasian | 1+ | <1 | TC | Non-pCR |
| 14 | 63 years | White or Caucasian | 1+ | <1 | KN-522(pembro-paclitaxel) | Non-pCR |
| 15 | 66 years | White or Caucasian | 1+ | 0 | pembro+AC; Sacituzumab | Non-pCR |
| 16 | 50 years | Other | 3+ | 0 | TCHP | pCR |
| 17 | 49 years | Black or African American | 2+ | <1 | AC Taxol | Non-pCR |
| 18 | 49 years | Other | 2+ | <10 | AC Taxol | pCR |
| 19 | 39 years | White or Caucasian | 1+ | <1 | Taxol | Non-pCR |
| 20 | 62 years | Black or African American | 1+ | <1 | AC Taxol | Non-pCR |
| 23 | 36 years | White or Caucasian | 1+ | 0 | AC X4 | Non-pCR |
| 24 | 66 years | White or Caucasian | 1+ | <10 | Carbo+Gem | Non-pCR |
| 26 | 36 years | White or Caucasian | 1+ | 0 | Pembro+Taxol+Carbo | Non-pCR |

**Supplementary Table 1:** Patient information including age, race, HER status, ER/PR status, treatment strategy and sensitivity to therapy [TCHP (Taxotere (docetaxel), Carboplatin, Herceptin (trastuzumab), and Perjeta (pertuzumab)); AC (Doxorubicin (Adriamycin) and Cyclophosphamide (Cytoxan)); pembro (pembrolizumab); TC (Taxotere (docetaxel) and Cytoxan (cyclophosphamide)); Carbo (Carboplatin); Gem (Gemcitabine); DD (Dose Dense)], pCR (Pathological complete response)

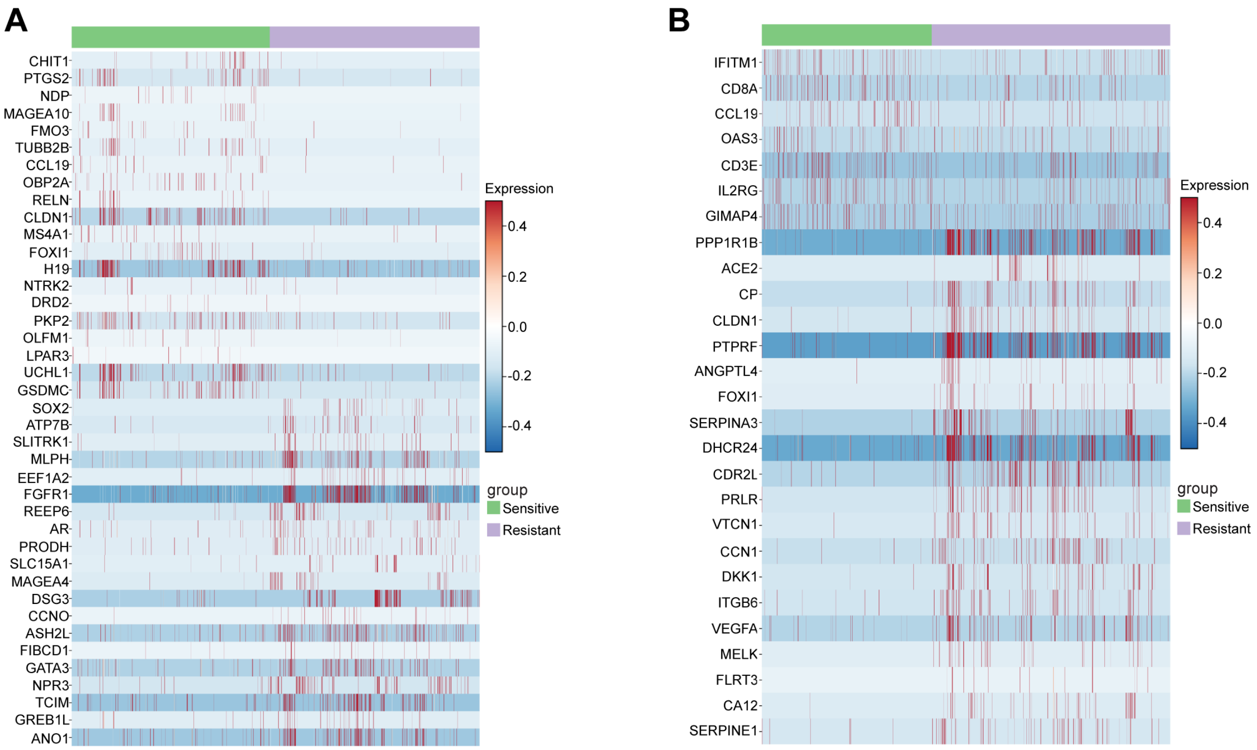


**Supplementary Figure 1.** Differential gene-expression programs distinguish sensitive and resistant HER2-low (**A**) and HER2-high tumors **(B).** Heatmaps show differentially expressed genes between sensitive and resistant tumors in the HER2-low and HER2-high groups.

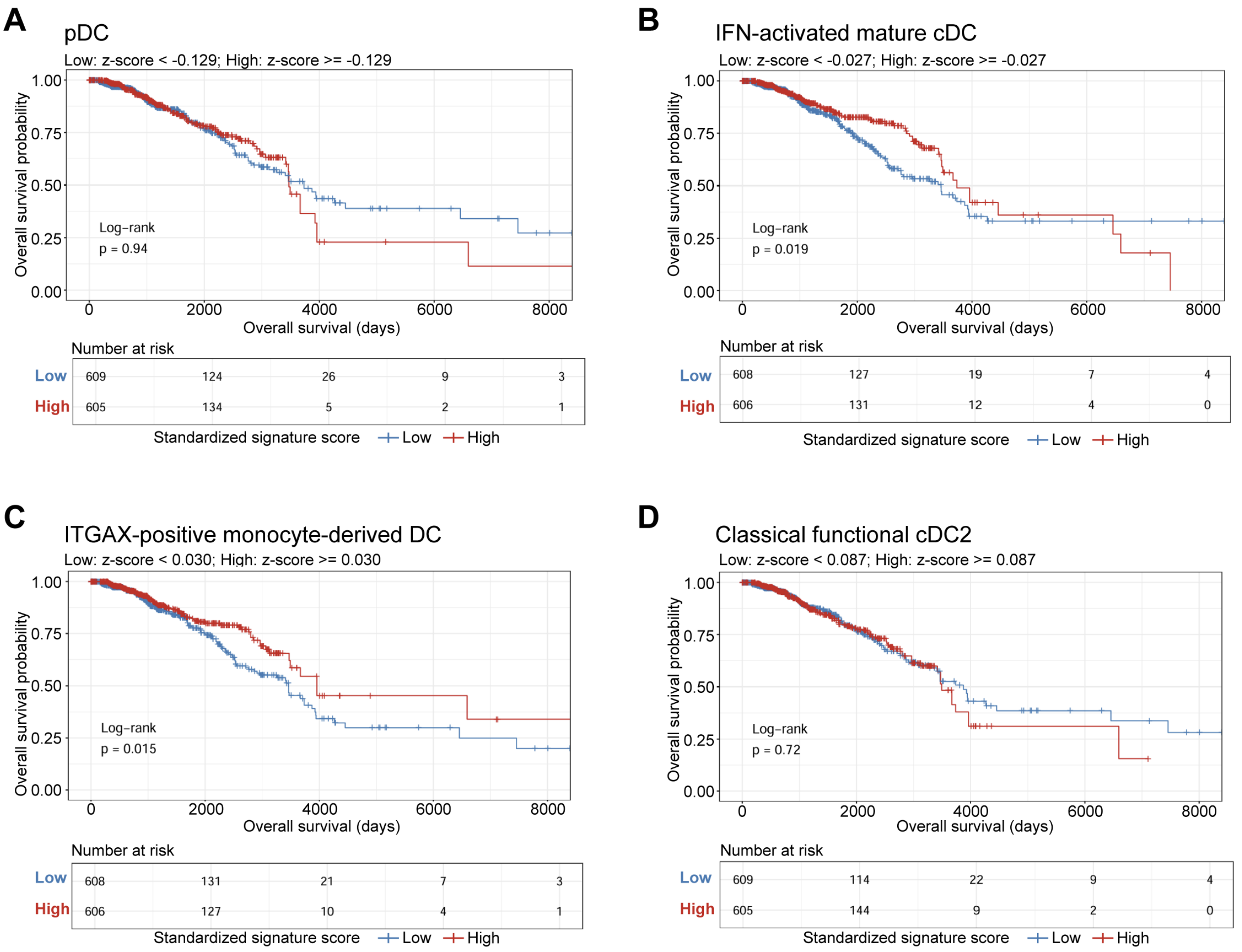


**Supplementary Figure 2.** Kaplan–Meier survival analysis of the TCGA-BRCA cohort (n = 1,214) stratified by dendritic-cell (DC) signature scores. Patients were dichotomized into high- and low-expression groups based on the median signature score derived from standardized (z-score) expression of marker genes for **(A)** plasmacytoid DC (CLEC4C, LILRA4, IL3RA) ((p = 0.94), **(B)** IFN-activated mature cDC (CXCL10, CXCL11, IFIT2) ((p = 0.019; FDR-adjusted p ≈ 0.032), **(C)** ITGAX-positive monocyte-derived DC (ITGAX) ((p = 0.015; FDR-adjusted p ≈ 0.032), and **(D)** classical functional cDC2 (P2RY13, CIITA) (p = 0.72). No significant survival differences were observed for the pDC or classical functional cDC2 signatures.

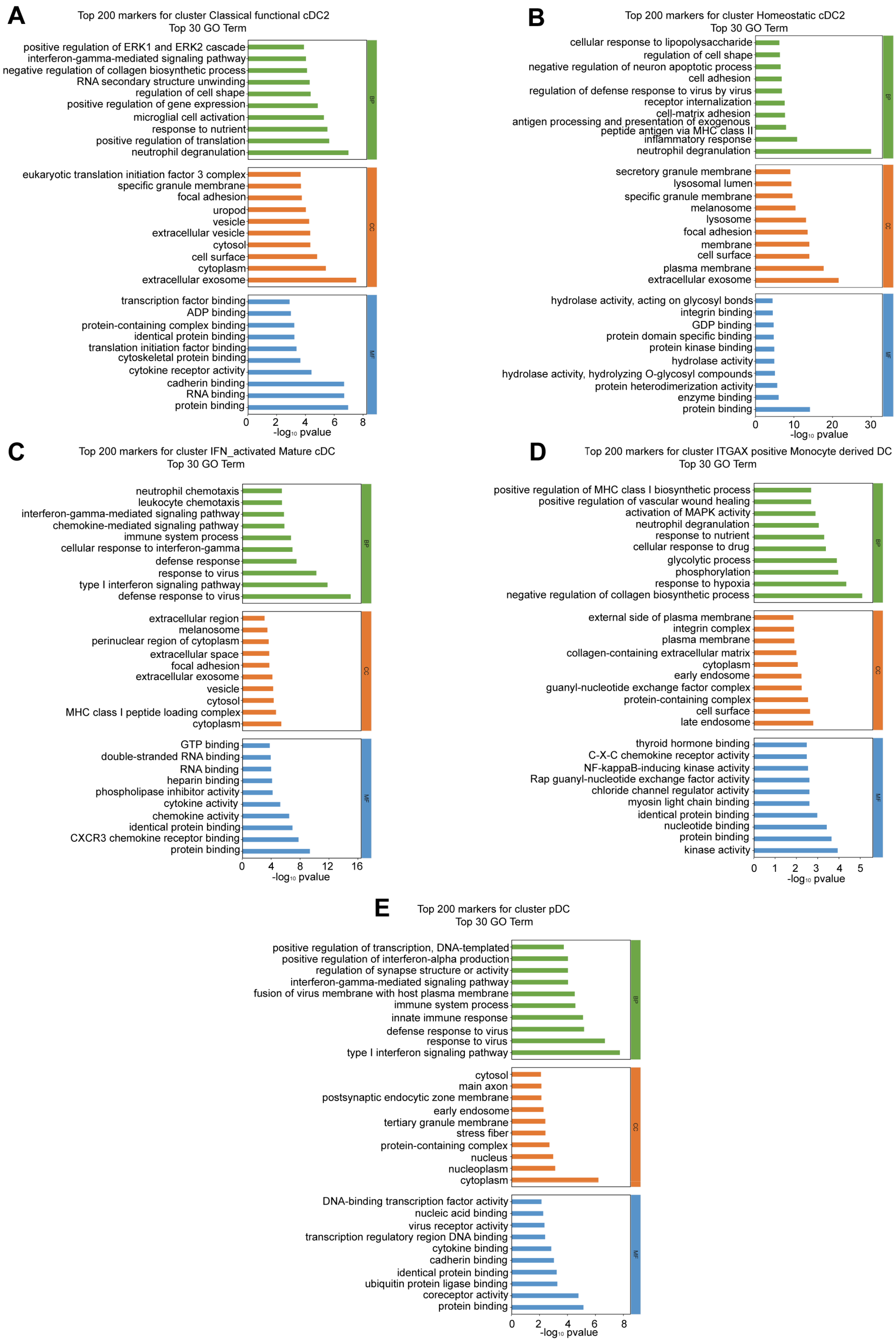

A
Top 200 markers for cluster Classical functional cDC2
Top 30 GO Term
positive regulation of ERK1 and ERK2 cascade
interferon-gamma-mediated signaling pathway
negative regulation of collagen biosynthetic process
RNA secondary structure unwinding
regulation of cell shape
positive regulation of gene expression
microglial cell activation
response to nutrient
positive regulation of translation
neutrophil degranulation
eukaryotic translation initiation factor 3 complex
specific granule membrane
focal adhesion
uropod
vesicle
extracellular vesicle
cytosol
cell surface
cytoplasm
extracellular exosome
transcription factor binding
ADP binding
protein-containing complex binding
identical protein binding
translation initiation factor binding
cytoskeletal protein binding
cytokine receptor activity
cadherin binding
RNA binding
protein binding
0 2 4 6 8
-log10 pvalue
BP
CC
MF
B
Top 200 markers for cluster Homeostatic cDC2
Top 30 GO Term
cellular response to lipopolysaccharide
regulation of cell shape
negative regulation of neuron apoptotic process
cell adhesion
regulation of defense response to virus by virus
receptor internalization
cell-matrix adhesion
antigen processing and presentation of exogenous peptide antigen via MHC class II
inflammatory response
neutrophil degranulation
secretory granule membrane
lysosomal lumen
specific granule membrane
melanosome
lysosome
focal adhesion
membrane
cell surface
plasma membrane
extracellular exosome
hydrolase activity, acting on glycosyl bonds
integrin binding
GDP binding
protein domain specific binding
protein kinase binding
hydrolase activity
hydrolase activity, hydrolyzing O-glycosyl compounds
protein heterodimerization activity
enzyme binding
protein binding
0 10 20 30
-log10 pvalue
BP
CC
MF
C
Top 200 markers for cluster IFN_activated Mature cDC
Top 30 GO Term
neutrophil chemotaxis
leukocyte chemotaxis
interferon-gamma-mediated signaling pathway
chemokine-mediated signaling pathway
immune system process
cellular response to interferon-gamma
defense response
response to virus
type I interferon signaling pathway
defense response to virus
extracellular region
melanosome
perinuclear region of cytoplasm
extracellular space
focal adhesion
extracellular exosome
vesicle
cytosol
MHC class I peptide loading complex
cytoplasm
GTP binding
double-stranded RNA binding
RNA binding
heparin binding
phospholipase inhibitor activity
cytokine activity
chemokine activity
identical protein binding
CXCR3 chemokine receptor binding
protein binding
0 4 8 12 16
-log10 pvalue
BP
CC
MF
D
Top 200 markers for cluster ITGAX positive Monocyte derived DC
Top 30 GO Term
positive regulation of MHC class I biosynthetic process
positive regulation of vascular wound healing
activation of MAPK activity
neutrophil degranulation
response to nutrient
cellular response to drug
glycolytic process
phosphorylation
response to hypoxia
negative regulation of collagen biosynthetic process
external side of plasma membrane
integrin complex
plasma membrane
collagen-containing extracellular matrix
cytoplasm
early endosome
guanyl-nucleotide exchange factor complex
protein-containing complex
cell surface
late endosome
thyroid hormone binding
C-X-C chemokine receptor activity
NF-kappaB-inducing kinase activity
Rap guanyl-nucleotide exchange factor activity
chloride channel regulator activity
myosin light chain binding
identical protein binding
nucleotide binding
protein binding
kinase activity
0 1 2 3 4 5
-log10 pvalue
BP
CC
MF
E
Top 200 markers for cluster pDC
Top 30 GO Term
positive regulation of transcription, DNA-templated
positive regulation of interferon-alpha production
regulation of synapse structure or activity
interferon-gamma-mediated signaling pathway
fusion of virus membrane with host plasma membrane
immune system process
innate immune response
defense response to virus
response to virus
type I interferon signaling pathway
cytosol
main axon
postsynaptic endocytic zone membrane
early endosome
tertiary granule membrane
stress fiber
protein-containing complex
nucleus
nucleoplasm
cytoplasm
DNA-binding transcription factor activity
nucleic acid binding
virus receptor activity
transcription regulatory region DNA binding
cytokine binding
cadherin binding
identical protein binding
ubiquitin protein ligase binding
coreceptor activity
protein binding
0 2 4 6 8
-log10 pvalue
BP
CC
MF

**Supplementary Figure 3. Dendritic cell-associated biological function.** Gene ontology showing biological processes associated with each DC subtype **(A)** classical functional cDC2 **(B)** homeostatic cDC2 **(C)** IFN-activated mature cDC **(D)** ITGAX-positive monocyte-derived DC **(E)** pDC

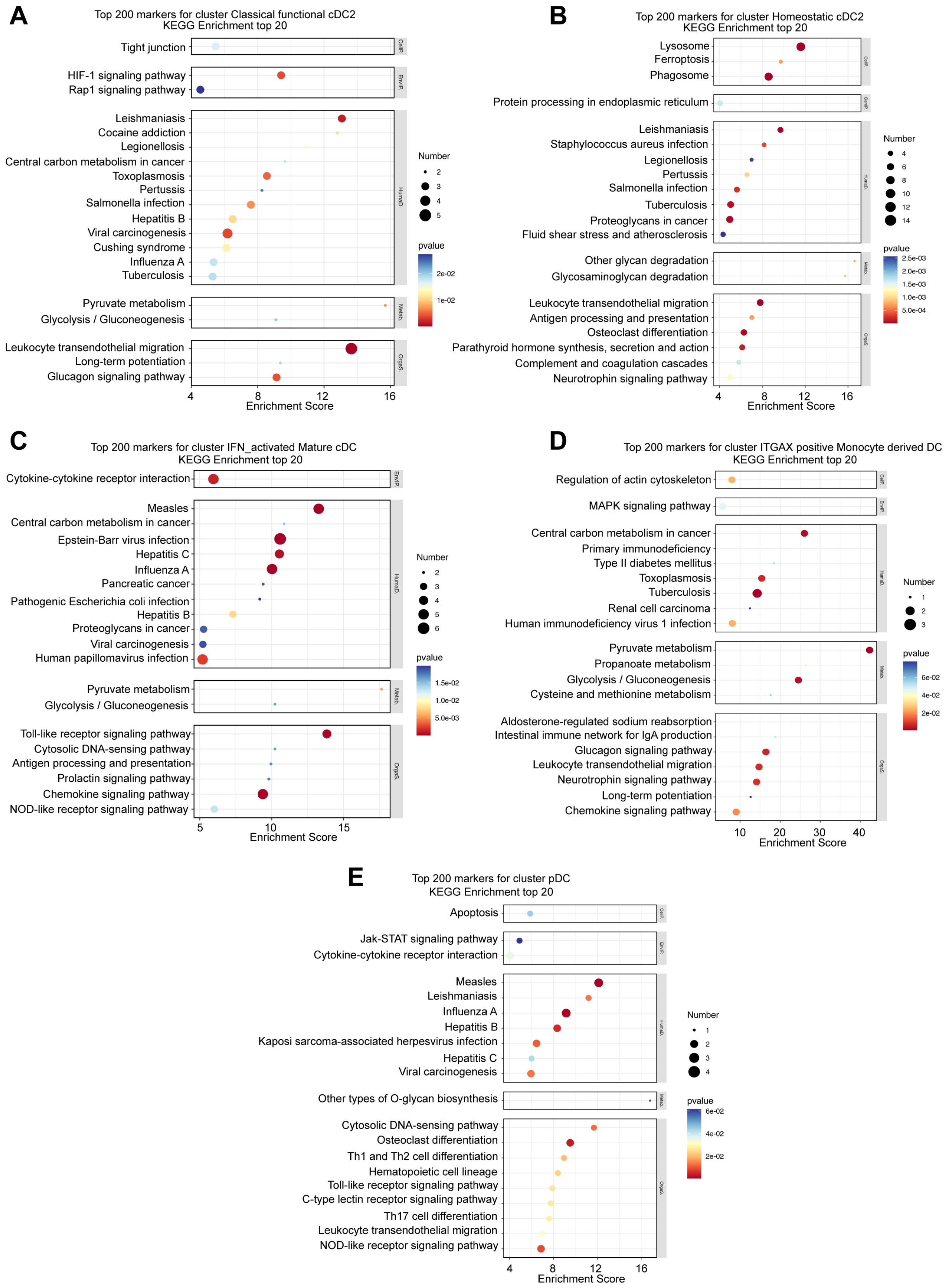

A
Top 200 markers for cluster Classical functional cDC2
KEGG Enrichment top 20
Tight junction
HIF-1 signaling pathway
Rap1 signaling pathway
Leishmaniasis
Cocaine addiction
Legionellosis
Central carbon metabolism in cancer
Toxoplasmosis
Pertussis
Salmonella infection
Hepatitis B
Viral carcinogenesis
Cushing syndrome
Influenza A
Tuberculosis
Pyruvate metabolism
Glycolysis / Gluconeogenesis
Leukocyte transendothelial migration
Long-term potentiation
Glucagon signaling pathway
4 8 12 16
Enrichment Score
Number
2 3 4 5
pvalue
2e-02
1e-02
B
Top 200 markers for cluster Homeostatic cDC2
KEGG Enrichment top 20
Lysosome
Ferroptosis
Phagosome
Protein processing in endoplasmic reticulum
Leishmaniasis
Staphylococcus aureus infection
Legionellosis
Pertussis
Salmonella infection
Tuberculosis
Proteoglycans in cancer
Fluid shear stress and atherosclerosis
Other glycan degradation
Glycosaminoglycan degradation
Leukocyte transendothelial migration
Antigen processing and presentation
Osteoclast differentiation
Parathyroid hormone synthesis, secretion and action
Complement and coagulation cascades
Neurotrophin signaling pathway
4 8 12 16
Enrichment Score
Number
4 6 8 10 12 14
pvalue
2.5e-03
2.0e-03
1.5e-03
1.0e-03
5.0e-04
C
Top 200 markers for cluster IFN_activated Mature cDC
KEGG Enrichment top 20
Cytokine-cytokine receptor interaction
Measles
Central carbon metabolism in cancer
Epstein-Barr virus infection
Hepatitis C
Influenza A
Pancreatic cancer
Pathogenic Escherichia coli infection
Hepatitis B
Proteoglycans in cancer
Viral carcinogenesis
Human papillomavirus infection
Pyruvate metabolism
Glycolysis / Gluconeogenesis
Toll-like receptor signaling pathway
Cytosolic DNA-sensing pathway
Antigen processing and presentation
Prolactin signaling pathway
Chemokine signaling pathway
NOD-like receptor signaling pathway
5 10 15
Enrichment Score
Number
2 3 4 5 6
pvalue
1.5e-02
1.0e-02
5.0e-03
D
Top 200 markers for cluster ITGAX positive Monocyte derived DC
KEGG Enrichment top 20
Regulation of actin cytoskeleton
MAPK signaling pathway
Central carbon metabolism in cancer
Primary immunodeficiency
Type II diabetes mellitus
Toxoplasmosis
Tuberculosis
Renal cell carcinoma
Human immunodeficiency virus 1 infection
Pyruvate metabolism
Propanoate metabolism
Glycolysis / Gluconeogenesis
Cysteine and methionine metabolism
Aldosterone-regulated sodium reabsorption
Intestinal immune network for IgA production
Glucagon signaling pathway
Leukocyte transendothelial migration
Neurotrophin signaling pathway
Long-term potentiation
Chemokine signaling pathway
10 20 30 40
Enrichment Score
Number
1 2 3
pvalue
6e-02
4e-02
2e-02
E
Top 200 markers for cluster pDC
KEGG Enrichment top 20
Apoptosis
Jak-STAT signaling pathway
Cytokine-cytokine receptor interaction
Measles
Leishmaniasis
Influenza A
Hepatitis B
Kaposi sarcoma-associated herpesvirus infection
Hepatitis C
Viral carcinogenesis
Other types of O-glycan biosynthesis
Cytosolic DNA-sensing pathway
Osteoclast differentiation
Th1 and Th2 cell differentiation
Hematopoietic cell lineage
Toll-like receptor signaling pathway
C-type lectin receptor signaling pathway
Th17 cell differentiation
Leukocyte transendothelial migration
NOD-like receptor signaling pathway
4 8 12 16
Enrichment Score
Number
1 2 3 4
pvalue
6e-02
4e-02
2e-02

**Supplementary Figure 4. Functional diversity of DC population.** KEGG pathway enrichment analysis shows heterogeneity in the function of the different subsets of DC **(A)** classical functional cDC2 **(B)** homeostatic cDC2 **(C)** IFN-activated mature cDC **(D)** ITGAX-positive monocyte-derived DC **(E)** pDC.

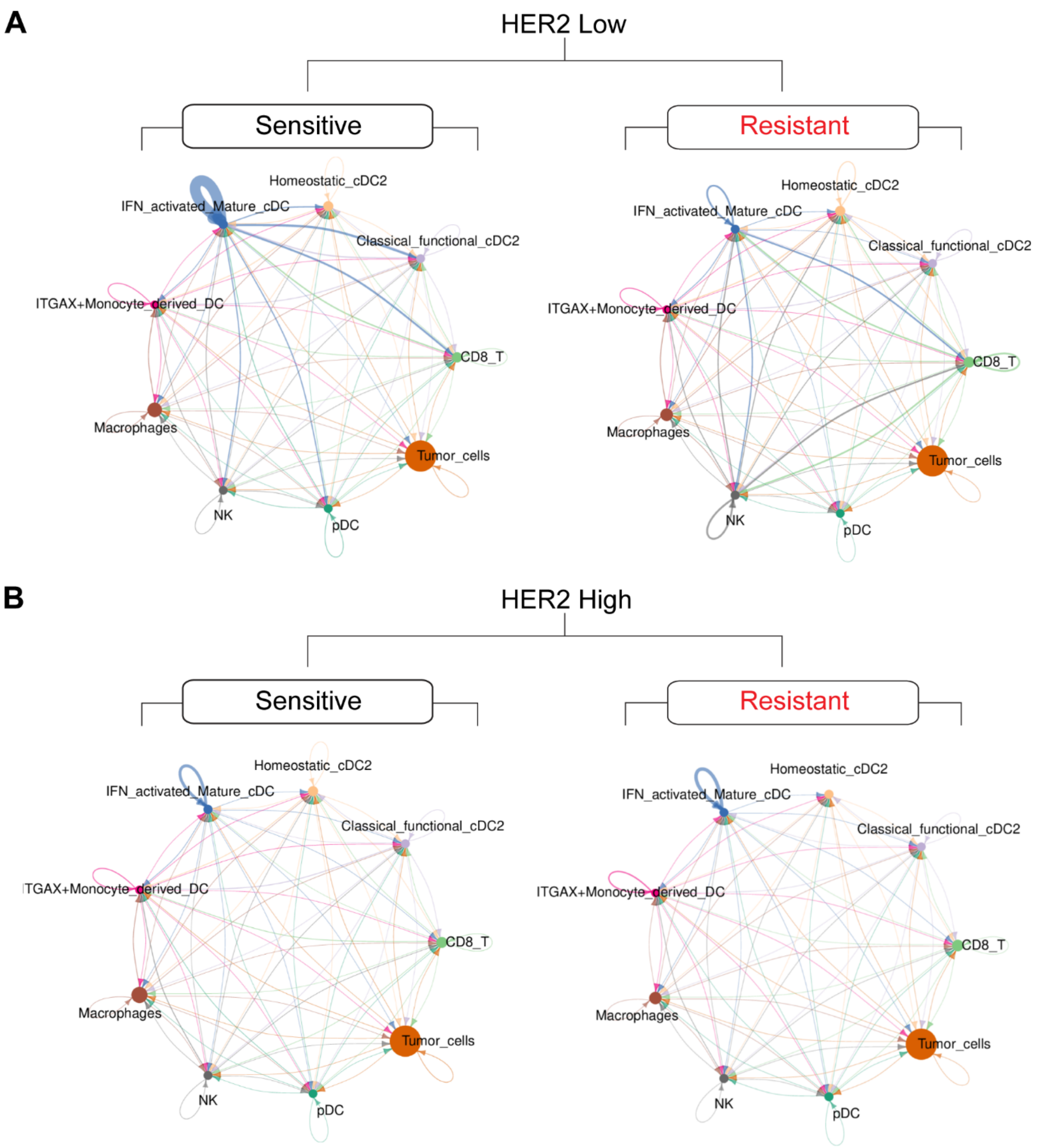


**Supplementary Figure 5. Cell-cell communication shows interaction strength in HER2-low and HER2-high breast tumors. (A**) Interaction strength in HER2-low sensitive vs. resistant tumors **(B)** Interaction strength in HER2-high sensitive vs. resistant tumors.

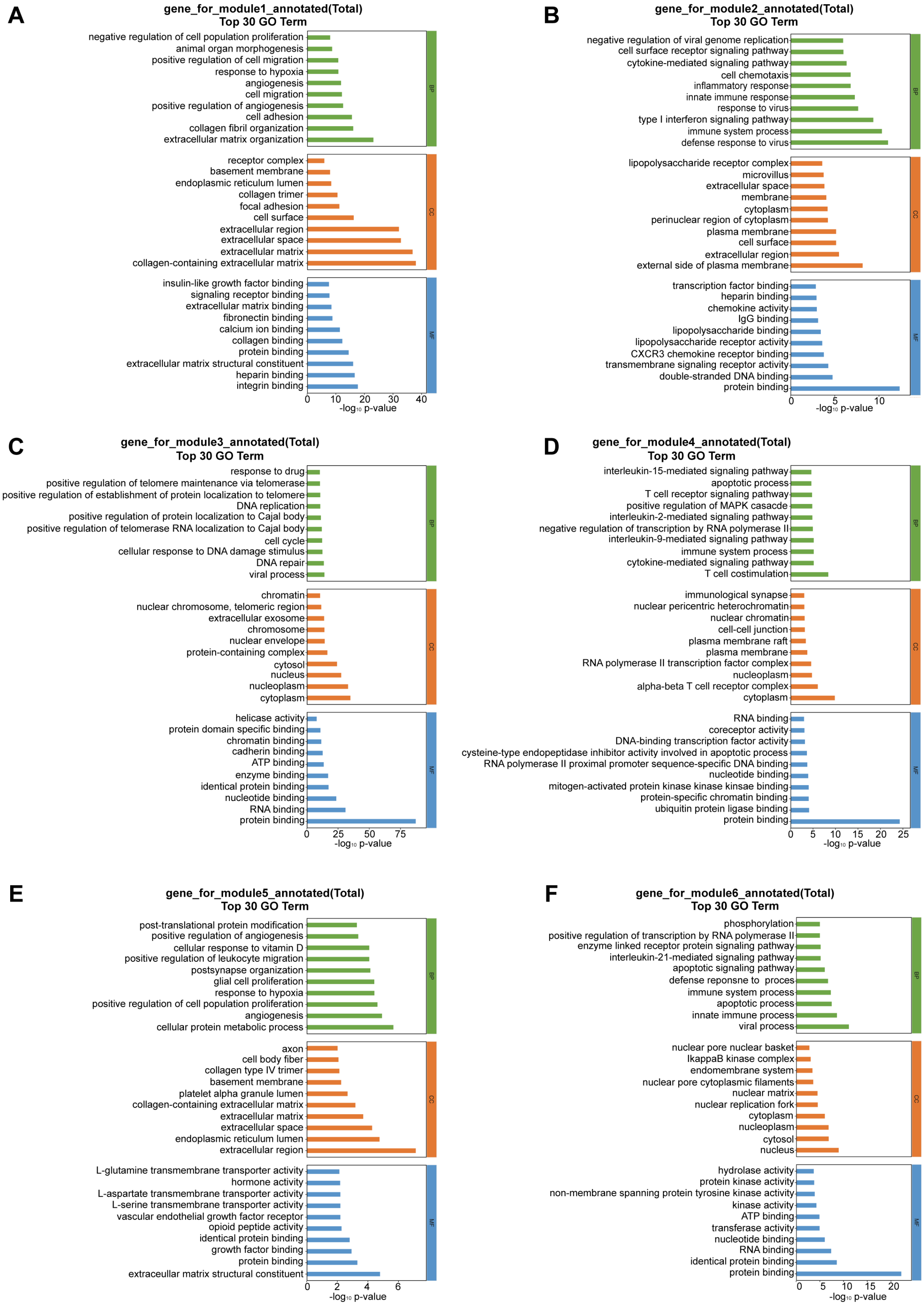

A
gene_for_module1_annotated(Total)
Top 30 GO Term
negative regulation of cell population proliferation
animal organ morphogenesis
positive regulation of cell migration
response to hypoxia
angiogenesis
cell migration
positive regulation of angiogenesis
cell adhesion
collagen fibril organization
extracellular matrix organization
BP
receptor complex
basement membrane
endoplasmic reticulum lumen
collagen trimer
focal adhesion
cell surface
extracellular region
extracellular space
extracellular matrix
collagen-containing extracellular matrix
CC
insulin-like growth factor binding
signaling receptor binding
extracellular matrix binding
fibronectin binding
calcium ion binding
collagen binding
protein binding
extracellular matrix structural constituent
heparin binding
integrin binding
MF
0
10
20
30
40
-log10 p-value
B
gene_for_module2_annotated(Total)
Top 30 GO Term
negative regulation of viral genome replication
cell surface receptor signaling pathway
cytokine-mediated signaling pathway
cell chemotaxis
inflammatory response
innate immune response
response to virus
type I interferon signaling pathway
immune system process
defense response to virus
BP
lipopolysaccharide receptor complex
microvillus
extracellular space
membrane
cytoplasm
perinuclear region of cytoplasm
plasma membrane
cell surface
extracellular region
external side of plasma membrane
CC
transcription factor binding
heparin binding
chemokine activity
IgG binding
lipopolysaccharide binding
lipopolysaccharide receptor activity
CXCR3 chemokine receptor binding
transmembrane signaling receptor activity
double-stranded DNA binding
protein binding
MF
0
5
10
-log10 p-value
C
gene_for_module3_annotated(Total)
Top 30 GO Term
response to drug
positive regulation of telomere maintenance via telomerase
positive regulation of establishment of protein localization to telomere
DNA replication
positive regulation of protein localization to Cajal body
positive regulation of telomerase RNA localization to Cajal body
cell cycle
cellular response to DNA damage stimulus
DNA repair
viral process
BP
chromatin
nuclear chromosome, telomeric region
extracellular exosome
chromosome
nuclear envelope
protein-containing complex
cytosol
nucleus
nucleoplasm
cytoplasm
CC
helicase activity
protein domain specific binding
chromatin binding
cadherin binding
ATP binding
enzyme binding
identical protein binding
nucleotide binding
RNA binding
protein binding
MF
0
25
50
75
-log10 p-value
D
gene_for_module4_annotated(Total)
Top 30 GO Term
interleukin-15-mediated signaling pathway
apoptotic process
T cell receptor signaling pathway
positive regulation of MAPK casacde
interleukin-2-mediated signaling pathway
negative regulation of transcription by RNA polymerase II
interleukin-9-mediated signaling pathway
immune system process
cytokine-mediated signaling pathway
T cell costimulation
BP
immunological synapse
nuclear pericentric heterochromatin
nuclear chromatin
cell-cell junction
plasma membrane raft
plasma membrane
RNA polymerase II transcription factor complex
nucleoplasm
alpha-beta T cell receptor complex
cytoplasm
CC
RNA binding
coreceptor activity
DNA-binding transcription factor activity
cysteine-type endopeptidase inhibitor activity involved in apoptotic process
RNA polymerase II proximal promoter sequence-specific DNA binding
nucleotide binding
mitogen-activated protein kinase kinase kinsae binding
protein-specific chromatin binding
ubiquitin protein ligase binding
protein binding
MF
0
5
10
15
20
25
-log10 p-value
E
gene_for_module5_annotated(Total)
Top 30 GO Term
post-translational protein modification
positive regulation of angiogenesis
cellular response to vitamin D
positive regulation of leukocyte migration
postsynapse organization
glial cell proliferation
response to hypoxia
positive regulation of cell population proliferation
angiogenesis
cellular protein metabolic process
BP
axon
cell body fiber
collagen type IV trimer
basement membrane
platelet alpha granule lumen
collagen-containing extracellular matrix
extracellular matrix
extracellular space
endoplasmic reticulum lumen
extracellular region
CC
L-glutamine transmembrane transporter activity
hormone activity
L-aspartate transmembrane transporter activity
L-serine transmembrane transporter activity
vascular endothelial growth factor receptor
opioid peptide activity
identical protein binding
growth factor binding
protein binding
extraceullar matrix structural constituent
MF
0
2
4
6
-log10 p-value
F
gene_for_module6_annotated(Total)
Top 30 GO Term
phosphorylation
positive regulation of transcription by RNA polymerase II
enzyme linked receptor protein signaling pathway
interleukin-21-mediated signaling pathway
apoptotic signaling pathway
defense reponsne to proces
immune system process
apoptotic process
innate immune process
viral process
BP
nuclear pore nuclear basket
IkappaB kinase complex
endomembrane system
nuclear pore cytoplasmic filaments
nuclear matrix
nuclear replication fork
cytoplasm
nucleoplasm
cytosol
nucleus
CC
hydrolase activity
protein kinase activity
non-membrane spanning protein tyrosine kinase activity
kinase activity
ATP binding
transferase activity
nucleotide binding
RNA binding
identical protein binding
protein binding
MF
0
5
10
15
20
-log10 p-value

**Supplementary Figure 6. Gene ontology enrichment of pseudotime-associated modules in HER2-low and HER2-high breast tumors.** Biological processes as it relates to pseudotime-associated genes were clustered into six expression modules **(A)** module 1 **(B)** module 2 **(C)** module 3 **(D)** module 4 **(E)** module 5 **(F)** module 6

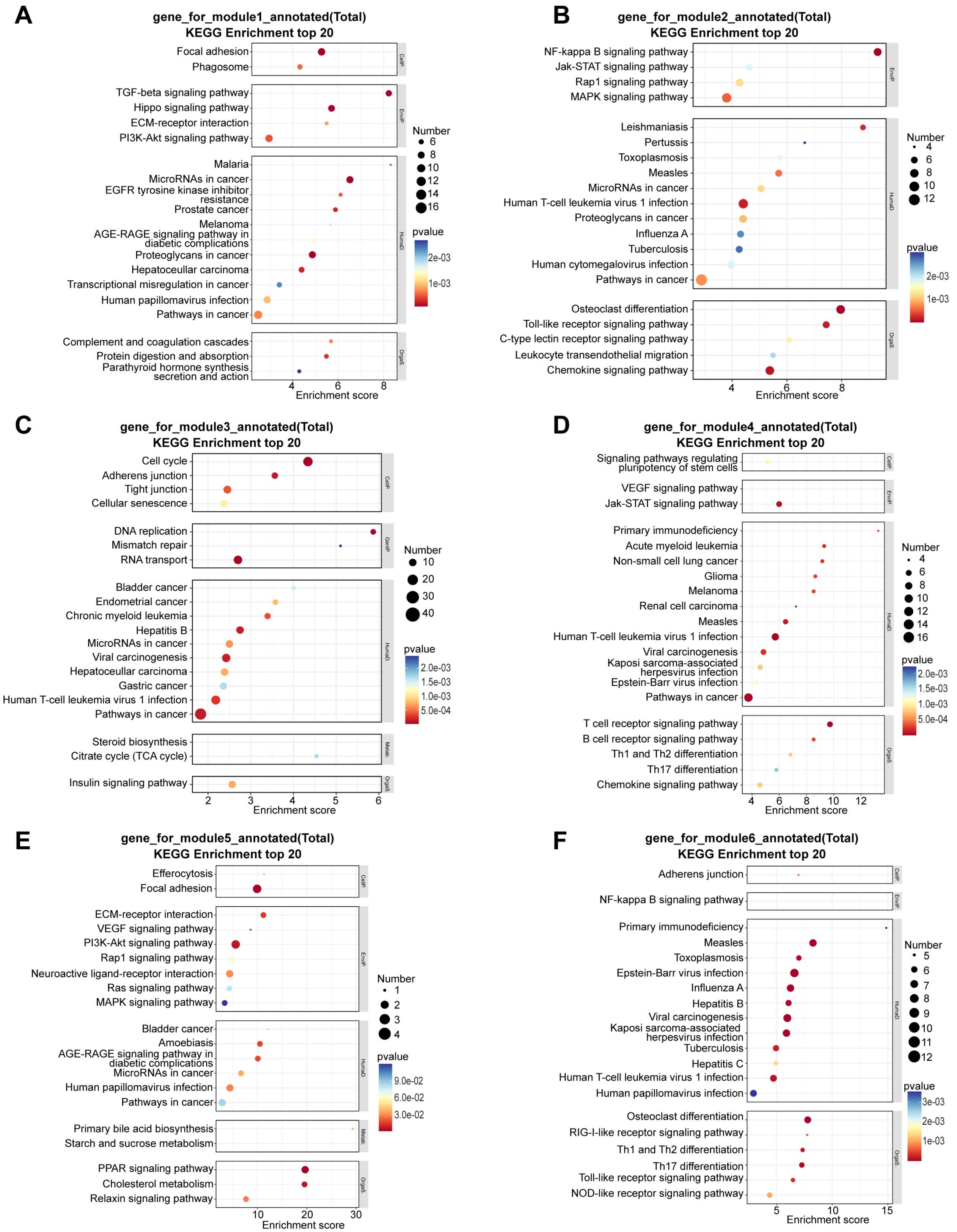
A
gene_for_module1_annotated(Total)
KEGG Enrichment top 20
Focal adhesion
Phagosome
TGF-beta signaling pathway
Hippo signaling pathway
ECM-receptor interaction
PI3K-Akt signaling pathway
Malaria
MicroRNAs in cancer
EGFR tyrosine kinase inhibitor resistance
Prostate cancer
Melanoma
AGE-RAGE signaling pathway in diabetic complications
Proteoglycans in cancer
Hepatocellular carcinoma
Transcriptional misregulation in cancer
Human papillomavirus infection
Pathways in cancer
Complement and coagulation cascades
Protein digestion and absorption
Parathyroid hormone synthesis secretion and action
4 6 8
Enrichment score
Number
6 8 10 12 14 16
pvalue
2e-03
1e-03
B
gene_for_module2_annotated(Total)
KEGG Enrichment top 20
NF-kappa B signaling pathway
Jak-STAT signaling pathway
Rap1 signaling pathway
MAPK signaling pathway
Leishmaniasis
Pertussis
Toxoplasmosis
Measles
MicroRNAs in cancer
Human T-cell leukemia virus 1 infection
Proteoglycans in cancer
Influenza A
Tuberculosis
Human cytomegalovirus infection
Pathways in cancer
Osteoclast differentiation
Toll-like receptor signaling pathway
C-type lectin receptor signaling pathway
Leukocyte transendothelial migration
Chemokine signaling pathway
4 6 8
Enrichment score
Number
4 6 8 10 12
pvalue
2e-03
1e-03
C
gene_for_module3_annotated(Total)
KEGG Enrichment top 20
Cell cycle
Adherens junction
Tight junction
Cellular senescence
DNA replication
Mismatch repair
RNA transport
Bladder cancer
Endometrial cancer
Chronic myeloid leukemia
Hepatitis B
MicroRNAs in cancer
Viral carcinogenesis
Hepatocellular carcinoma
Gastric cancer
Human T-cell leukemia virus 1 infection
Pathways in cancer
Steroid biosynthesis
Citrate cycle (TCA cycle)
Insulin signaling pathway
2 3 4 5 6
Enrichment score
Number
10 20 30 40
pvalue
2.0e-03
1.5e-03
1.0e-03
5.0e-04
D
gene_for_module4_annotated(Total)
KEGG Enrichment top 20
Signaling pathways regulating pluripotency of stem cells
VEGF signaling pathway
Jak-STAT signaling pathway
Primary immunodeficiency
Acute myeloid leukemia
Non-small cell lung cancer
Glioma
Melanoma
Renal cell carcinoma
Measles
Human T-cell leukemia virus 1 infection
Viral carcinogenesis
Kaposi sarcoma-associated herpesvirus infection
Epstein-Barr virus infection
Pathways in cancer
T cell receptor signaling pathway
B cell receptor signaling pathway
Th1 and Th2 differentiation
Th17 differentiation
Chemokine signaling pathway
4 6 8 10 12
Enrichment score
Number
4 6 8 10 12 14 16
pvalue
2.0e-03
1.5e-03
1.0e-03
5.0e-04
E
gene_for_module5_annotated(Total)
KEGG Enrichment top 20
Efferocytosis
Focal adhesion
ECM-receptor interaction
VEGF signaling pathway
PI3K-Akt signaling pathway
Rap1 signaling pathway
Neuroactive ligand-receptor interaction
Ras signaling pathway
MAPK signaling pathway
Bladder cancer
Amoebiasis
AGE-RAGE signaling pathway in diabetic complications
MicroRNAs in cancer
Human papillomavirus infection
Pathways in cancer
Primary bile acid biosynthesis
Starch and sucrose metabolism
PPAR signaling pathway
Cholesterol metabolism
Relaxin signaling pathway
10 20 30
Enrichment score
Number
1 2 3 4
pvalue
9.0e-02
6.0e-02
3.0e-02
F
gene_for_module6_annotated(Total)
KEGG Enrichment top 20
Adherens junction
NF-kappa B signaling pathway
Primary immunodeficiency
Measles
Toxoplasmosis
Epstein-Barr virus infection
Influenza A
Hepatitis B
Viral carcinogenesis
Kaposi sarcoma-associated herpesvirus infection
Tuberculosis
Hepatitis C
Human T-cell leukemia virus 1 infection
Human papillomavirus infection
Osteoclast differentiation
RIG-I-like receptor signaling pathway
Th1 and Th2 differentiation
Th17 differentiation
Toll-like receptor signaling pathway
NOD-like receptor signaling pathway
5 10 15
Enrichment score
Number
5 6 7 8 9 10 11 12
pvalue
3e-03
2e-03
1e-03

**Supplementary Figure 7. Functional diversity of pseudotime-associated genes in HER2-low and HER2-high breast tumors.** Spatially resolved expression profiles ordered along a pseudotime trajectory across 6 modules **(A)** module 1 **(B)** module 2 **(C)** module 3 **(D)** module 4 **(E)** module 5 **(F)** module 6
